# An affinity-matured DLL4 ligand for broad-spectrum activation and inhibition of Notch signaling

**DOI:** 10.1101/2022.03.07.483330

**Authors:** David Gonzalez-Perez, Satyajit Das, Elliot Medina, Daniel Antfolk, Emily D. Egan, Stephen C. Blacklow, Paulo C. Rodriguez, Vincent C. Luca

## Abstract

The Notch pathway regulates cell fate decisions and is an emerging target for regenerative and cancer therapies. Recombinant Notch ligands are attractive candidates for modulating Notch signaling; however, their intrinsically low receptor-binding affinity restricts their utility in biomedical applications. To overcome this limitation, we evolved variants of the ligand Delta-like 4 (DLL4) with enhanced affinity and cross-reactivity. A consensus variant with maximized binding affinity, Delta^MAX^, engages human and murine Notch receptors with 500- to 1000-fold increased affinity compared to wild-type human DLL4. Delta^MAX^ also potently activates human Notch in plate-bound, bead-bound, and cellular formats. When administered as a soluble decoy, Delta^MAX^ inhibits Notch activation in response to either Delta-like (DLL) or Jagged (Jag) ligands, highlighting its utility as both an agonist and antagonist. Finally, we demonstrate that Delta^MAX^ stimulates increased proliferation and expression of effector mediators in primary activated human T cells. Taken together, our data defines Delta^MAX^ as a versatile biotechnological tool for broad-spectrum activation or inhibition of Notch signaling.

## INTRODUCTION

The Notch pathway is a conserved signaling system that regulates metazoan development and cell fate decisions. In mammals, the core Notch signaling network consists of four Notch receptor paralogs (Notch1-4) and the activating ligands Delta-like 1 (DLL1), Delta-like 4 (DLL4), Jagged1 (Jag1), and Jagged2 (Jag2). Notch signaling occurs when cells expressing Notch proteins (signal receivers) interact with adjacent cells expressing DLL or Jag ligands (signal senders)^1–4^. Following ligand engagement, endocytosis of the DLL or Jag protein into the sender cell exerts a “pulling” force that destabilizes the negative regulatory region (NRR) of the Notch extracellular domain (ECD)^5, 6^. This mechanical tension exposes a proteolytic cleavage site (S2) that is processed by the ADAM10 metalloprotease^7, 8^. ADAM10 cleavage results in the shedding of the Notch ECD, which in turn sensitizes the Notch transmembrane domain to cleavage by the intramembrane protease gamma-secretase^1, 9, 10^. This second proteolytic event releases the Notch intracellular domain (NICD) from the plasma membrane and it translocates to the nucleus to serve as a transcriptional cofactor^1, 2, 11^.

Notch extracellular domains (ECDs) are comprised of 29 to 36 epidermal growth factor (EGF) domains and the juxtamembrane negative regulatory region (NRR). Ligand ECDs also have a modular structure and contain an N-terminal C2-like (MNNL) domain, a Delta/Serrate/Lag3 (DSL) domain, 6 to 16 EGF domains, and a cysteine rich domain that is only present in Jag proteins. Through structural, biochemical, and cell-based studies, we and others have determined that the C2, DSL, and EGF1-3 region of DLL or Jag engage five centrally positioned Notch EGF domains (EGF8-12 in Notch1) to initiate signaling^4, 12–15^. Interactions between Notch and ligand ECDs are low affinity^4, 13, 16^, and co-crystallization of rat DLL4:Notch1 and Jag1:Notch1 complexes required the incorporation of affinity-enhancing mutations into the ligands^14, 15^. Despite the low affinity of receptor for ligand, Notch signaling occurs productively *in vivo*, because receptor-ligand interactions may be strengthened by catch bond formation or multivalent binding^15, 17–19^.

Although the Notch transcriptional machinery is broadly conserved, preferential interactions have been described between receptor and ligand subtypes^1^. For example, DLL4 is a higher affinity ligand for Notch1 than is DLL1^13^, while DLL1 activates Notch1 and Notch2 roughly equivalently^16^. Signaling through specific Notch-ligand pairs is also associated with distinct functional outcomes. During angiogenesis, Notch1:DLL4 interactions inhibit tip sprouting and Notch1:Jag1 interactions promote vessel growth^20^, and Jagged2:Notch3 interactions stimulate the differentiation of γ/δ T cells^21^. Comparative studies have begun to identify differences in the structures^15^, binding affinities^13^, and signaling dynamics^19^ of selected ligands that may contribute to their unique functional properties. However, the precise molecular mechanisms controlling receptor- and ligand-specific signaling remain unclear.

Depending on cellular context, Notch signaling may induce the differentiation or proliferation of stem cells, and the broad influence of Notch on stem cell behavior has made the pathway an attractive target for regenerative therapeutics. Moreover, Notch signaling has been shown to have context-dependent tumor suppressor or oncogenic functions in human cancers^22^. The pleiotropic effects of Notch signaling suggest that both agonists and antagonists of Notch signaling will be valuable for biomedical applications. Several antagonists are currently in clinical trials for the treatment of cancer, including gamma-secretase inhibitors and monoclonal antibodies targeting individual Notch ECDs ^23–25^. On the other hand, the development of Notch agonists has been challenging because soluble drugs cannot exert the mechanical force necessary for receptor activation. Thus far, a single agonist antibody has been described for Notch3, which may be uniquely susceptible to antibody-mediated activation due to the inherent instability of the Notch3 NRR^26, 27^.

Recombinant DLL or Jag ECDs are attractive “one-size-fits-all” candidates for activating or inhibiting Notch signaling. In their soluble form, ligand ECDs act as untethered decoys that bind to Notch receptors and inhibit ligand-mediated activation. The on-target specificity of ligand ECDs may also be advantageous given that gamma-secretase cleaves a myriad of cellular substrates that are unrelated to the Notch pathway^28^. Alternatively, surface- or matrix-tethered DLL or Jag ligands stimulate Notch signaling, presumably because immobilization provides sufficient resistance force for receptor activation. Despite their potential to function as universal modulators of the Notch pathway, the intrinsically low affinities of DLL and Jag ligands limits their practical utility as biomedical tools. Additionally, the preferential binding observed between certain receptor:ligand pairs may restrict the activity of a given ligand to a subset of Notch paralogs.

To overcome the biochemical limitations of endogenous Notch ligands, we used yeast display and mutational engraftment to engineer potent, broadly-reactive human DLL4 variants. By combining mutations from multiple directed evolution campaigns, we developed a consensus variant, named Delta^MAX^, that binds with greatly enhanced affinity to human and murine Notch receptor paralogs. We determined that Delta^MAX^ is a more potent Notch activator than WT DLL4 in several commonly used ligand-presentation formats, and that the proliferation and expression of effector mediators in primary T cells is enhanced following conditioning stimulation with Delta^MAX^. Furthermore, the soluble Delta^MAX^ ECD functions as a potent, “pan-Notch” inhibitor by competing for the Notch ligand-binding site. Our collective insights demonstrate that affinity-matured Notch ligands are highly-effective, multifunctional tools for modulating Notch signaling.

## RESULTS

### Structure-guided engineering of high-affinity DLL4 variants

We employed a structure-guided engineering strategy to evolve broadly-reactive, high-affinity DLL4 variants. Crystal structures of DLL4:Notch1 and Jag1:Notch1 complexes revealed that the ligand C2 and DSL domains engage the Notch1 EGF12 and EGF11 domains, respectively. In each of these structures, the C2:EGF12 interface was named Site 1 and the DSL:EGF11 interface was named Site 2. However, an additional “Site 3” binding interface was also visualized in the Jag1:Notch1 structure and forms between Jag1 EGF1-3 and Notch1 EGF8-10^14, 15^. Solution binding studies demonstrated that Site 3 contributes substantially to Jag1:Notch1 interactions, but has a minimal effect on DLL4:Notch1 interactions^15^. Therefore, we designed a site-directed mutant library to select for DLL4 mutations that recapitulate the Site 3 interaction observed in Notch1:Jag1. The library was generated by varying nine DLL4 residues (H256, N257, T271, L279, F280, T289, S301, N302, Q305) that were analogous to Site 3 interface residues in Jag1 EGF2 and EGF3. We employed a “light mutagenesis” approach in which each interface residue was allowed to encode for the WT DLL4 residue, the equivalent Jag1 residue, or biochemically similar residues (Fig. 1a, 1b).

**Fig 1.**
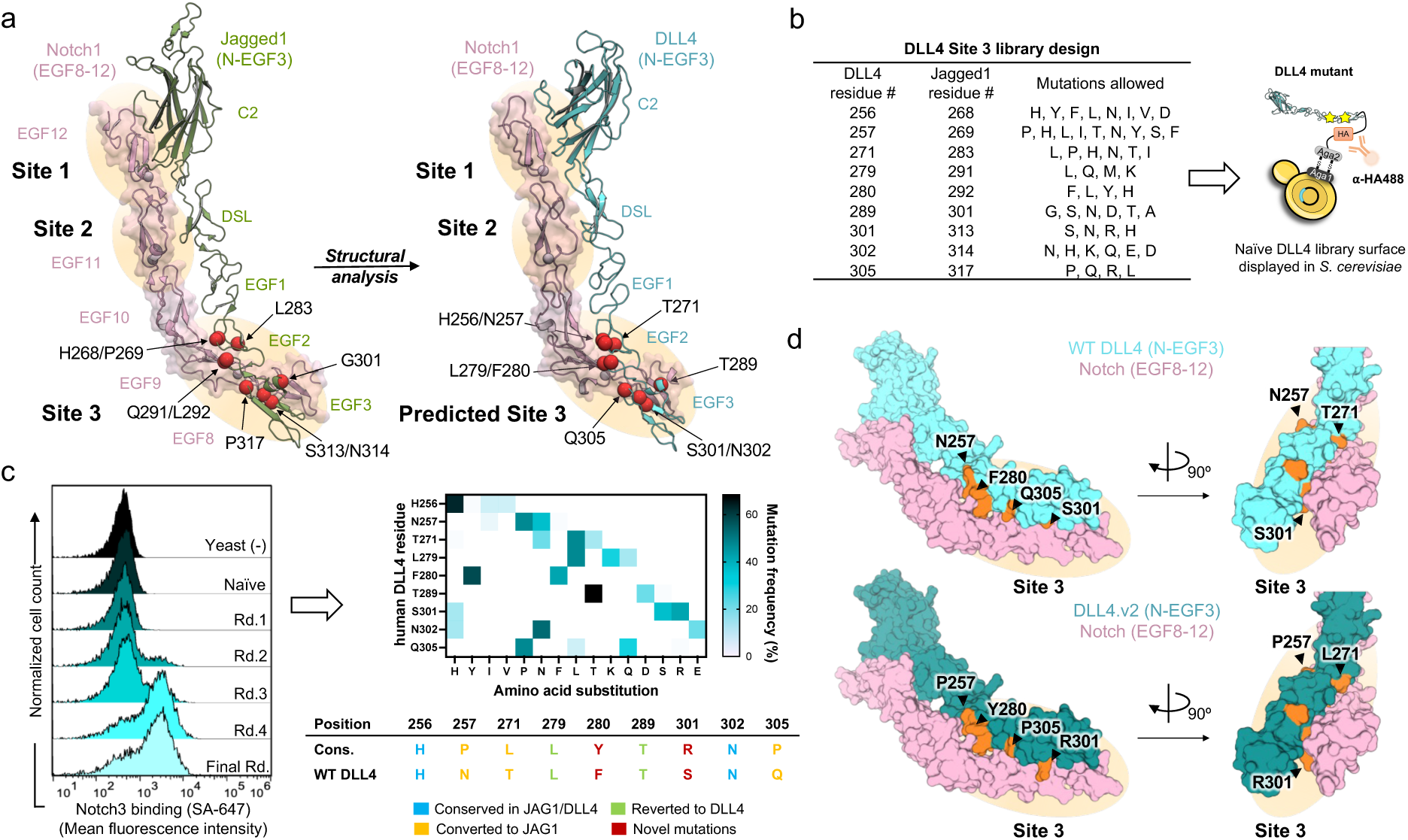
Yeast display selection of affinity-enhancing mutations in DLL4. (**A**) Cartoon schematic of the design strategy for the DLL4 Site 3 mutant library. Red spheres depict mutated interface residues. (**B**) Table depicting DLL4 interface residue positions and the mutations allowed at each position and schematic of DLL4 yeast display construct. Yellow stars indicate mutations. (**C**) Flow cytometry histogram plots of yeast stained with fluorescently-labeled Notch3 protein following each round of selection (left), and a table indicating the frequency of mutations and consensus DLL4 mutant sequence (right). (**D**) Structural models highlighting the mutated residues in DLL4.v2. WT residues in DLL4 and mutated residues in DLL4.v2 are colored orange and shown in surface representation. DLL4 models were based on DLL4:Notch1 (PDB ID: 4XL1, for C2-DSL-EGF1 domains) and Jagged1:Notch1 (PDB ID:5UK5, for EGF1-EGF3) structures.

We used yeast display to select high-affinity Notch binders from our DLL4 mutant library (Fig. 1b). The library was stained with Notch1 or Notch3 ECD constructs containing the ligand-binding domains (Notch1 EGF8-12, Notch3 EGF5-12) and several rounds of selections were performed to isolate high-affinity binders. The Notch3-selected yeast had the greatest enrichment of Notch-binders, and sequencing of individual colonies revealed twenty-one different mutations at interface positions (Fig. 1c, Extended Data Fig. 1). A clone containing five mutations (N257P, T271L, F280Y, S301R, and Q305P) was overrepresented in the samples and was selected for further characterization. This clone is henceforth referred to as DLL4.v2. Analysis of the DLL4.v2 sequence revealed that F280Y and S301R are novel mutations that are not present in other mammalian Notch ligands (Extended Data Fig. 2), and that N257P, T271L, and Q305P had converted to the Jag1 residues. The other four mutated positions (DLL4 residues H256, L279, T289, and N302) reverted to the wild-type sequence (Fig. 1c). Notably, N257P was previously introduced into DLL4 to recreate the “DOS motif” found in Jag1^2, 29^, and this substitution increased receptor binding and signaling^30^.

We analyzed the DLL4.v2 mutations in the context of Site 3 of the Notch1:Jag1 complex structure to gain insight into their mechanisms of affinity-enhancement (Fig. 1d, Extended Data Fig. 3). Based on this analysis, we predict that the N257P, T271L, and Q305P mutations enhance binding by improving hydrophobic packing at the binding interface (Extended Data Fig. 3a, 3b, 3c). The F280Y mutation may stabilize the overall fold of the DLL4 protein by replacing the exposed hydrophobic Phe^280^ phenyl group with a more hydrophilic tyrosyl group (Extended Data Fig. 3d). Lastly, we predict that S301R enhances binding by introducing contacts between the guanidinium group of DLL4 Arg^301^ and the main-chain carbonyl of the Notch1 Cys^321^ and/or the aliphatic side chain of Val^322^ (Extended Data Fig. 3e).

### Generation of a high-affinity DLL4 consensus variant

We combined multiple sets of affinity-enhancing mutations in an attempt to engineer a DLL4 protein with maximal receptor-binding affinity. We used the high-affinity DLL4.v2 variant as a starting point for our construct design. We then engrafted five affinity-enhancing mutations (G28S, F107L, N118I, H194Y, and L206P) from our previously reported rat DLL4 variant (E12)^14^ onto the human DLL4.v2 scaffold (Fig. 2a). As the E12 mutations are located within the C2 and DSL domains (Site 1 and Site 2) of DLL4, we hypothesized that they would have an additive or synergistic effect when combined with the Site 3 mutations of DLL4.v2. The resulting consensus variant, Delta^MAX^, contains 10 total mutations: G28S, F107L, N118I, H194Y, L206P, N257P, T271L, F280Y, S301R, and Q305P (Fig. 2a).

**Fig 2.**
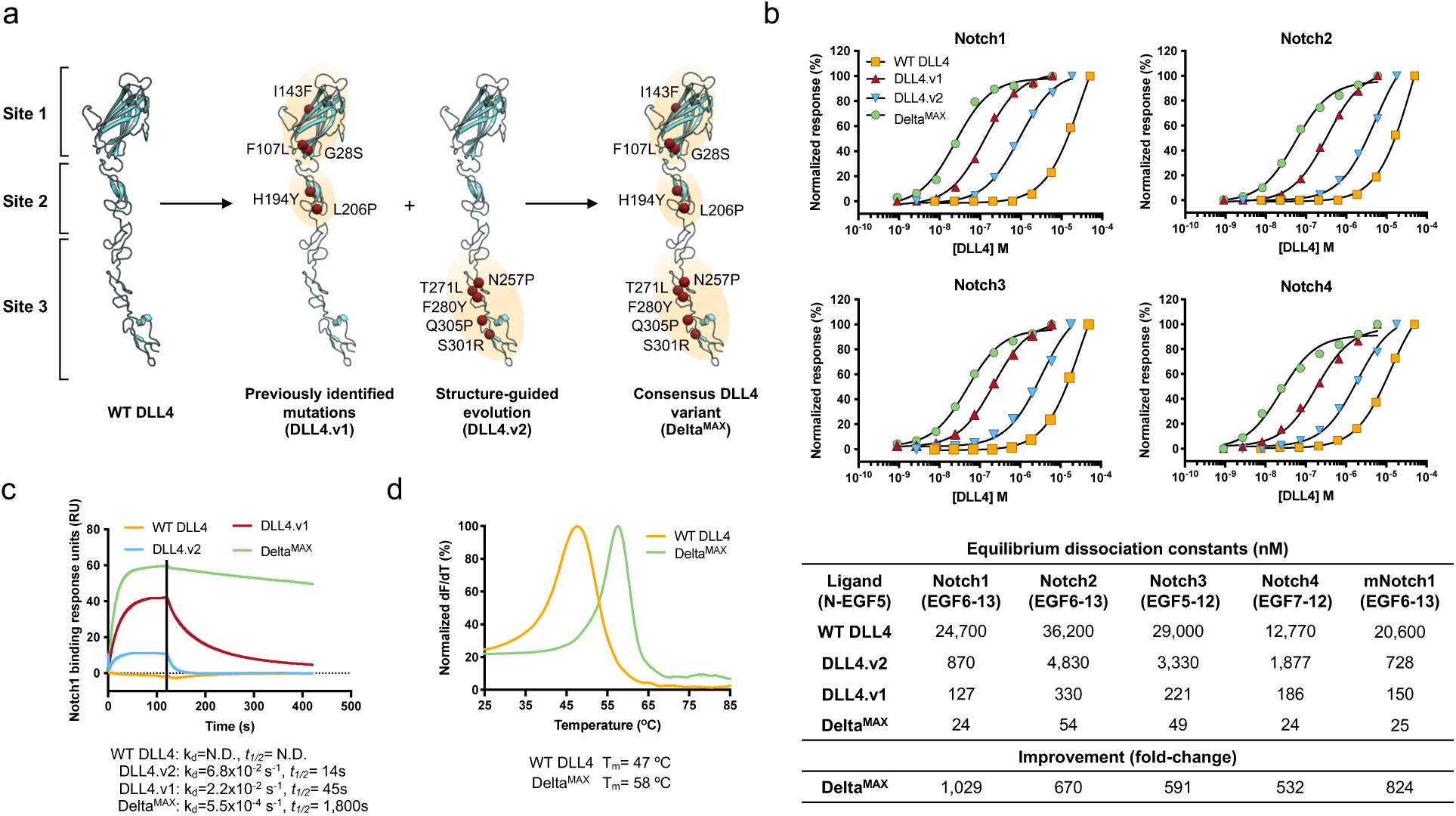
Biophysical characterization of Delta^MAX^ variant. (**A**) Schematic depicting the generation of DLL4^MAX^ through engraftment of DLL4.v1 and DLL4.v2 mutations. Red spheres indicate affinity-enhancing mutations. (**B**) SPR binding isotherms measuring DLL4 variant interactions with the ligand binding domains of Notch1-4. The table shows the K_D_ determined for each interaction and the fold-enhancement of DLL4^MAX^ affinity relative to WT DLL4. (**C**) Representative SPR sensograms depicting the binding of 200 nM concentrations of WT DLL4, DLL4.v1, DLL4.v2, and DLL4^MAX^ to Notch1 EGF6-13. (**D**) DSF was used to determine the T_m_ of WT DLL4 and Delta^MAX^.

### DLL4 variants bind to human and mouse Notch receptors with high-affinity

We used surface plasmon resonance (SPR) to determine the binding affinity between Delta^MAX^ and the ligand-binding regions of Notch1-4. As a basis for comparison, we also generated a WT DLL4 construct, a DLL4.v2 construct, and a “DLL4.v1” construct containing the five E12 mutations (G28S, F107L, N118I, H194Y, and L206P) (Fig. 2a, Extended Data Fig. 4). Analysis of the SPR data revealed that DLL4.v1, DLL4.v2, and Delta^MAX^ bound to all four human Notch receptors with enhanced affinity compared to WT DLL4 (Fig. 2b). DLL4.v1 bound to Notch1-4 with 70 to 200-fold increased affinity depending on the receptor subtype, and DLL4.v2 bound to Notch1-4 with 6 to 28-fold higher affinity. The larger contribution of Site 1 and Site 2 mutations from E12 to the interaction is consistent with previous data indicating that the C2 and DSL domains form the dominant receptor-binding interface. Delta^MAX^ bound to Notch1 with a dissociation constant (K_D_) of 24 nM, Notch2 with a K_D_ of 54 nM, Notch3 with a K_D_ of 49 nM, and Notch4 with a K_D_ of 24 nM. Compared to WT DLL4, the nanomolar affinity interactions between Delta^MAX^ and Notch1-4 represent enhancements of 1,100-fold, 670-fold, 591-fold, and 532-fold, respectively.

In addition to human Notch1-4, we measured the binding affinity between each DLL4 variant and the EGF6-13 region of murine Notch1 (mNotch1 EGF6-13) to test for cross-species reactivity (Fig. 2b, Extended Data Fig. 4, 5). Each ligand bound to murine Notch1 and human Notch1 with comparable affinities, and Delta^MAX^ bound to mNotch1 EGF6-13 with a K_D_ of 24 nM, which is nearly identical to its affinity for human Notch1 (24 nM K_D_). We attempted to fit the SPR data to compare the kinetic binding parameters between Notch and each of the four DLL4 constructs. The rapid dissociation of WT DLL4 precluded kinetic fitting of the binding data, however, visual inspection of the SPR sensograms indicates that the increasing affinities of DLL4.v1, DLL4.v2, and Delta^MAX^ are linked to progressive decreases in off-rate (Fig. 2c).

### Delta^MAX^ has improved expression and thermostability

We also tested whether the affinity-enhancing mutations impacted the expression and stability of the Delta^MAX^ protein. We performed a thermal denaturation experiment using differential scanning fluorimetry (DSF) to compare the melting temperatures of WT DLL4 and Delta^MAX^. We determined that the melting temperature (T_m_) of WT DLL4 was 47 °C and that the T_m_ of Delta^MAX^ was increased by 11 °C to 58 °C (Fig. 2d). Furthermore, expression yield of the Delta^MAX^ protein was increased by 2-fold compared to WT DLL4 (Extended Data Fig. 4g). Taken together, our SPR and DSF measurements indicate that Delta^MAX^ exhibits enhanced affinity, stability and expression while maintaining a broad reactivity profile against human and murine Notch receptors.

### Delta^MAX^ activates Notch more potently than WT DLL4

We performed a series of signaling assays to evaluate the Notch activation potency, signaling kinetics, and cross-reactivity of our high-affinity Delta^MAX^ ligand. We first characterized the signaling potency of Delta^MAX^ protein using fluorescent (H2B-Citrine) Notch1 reporter cells (Extended Data Fig. 6a, 7a)^31^. As our goal was to develop an accessible tool for modulating Notch activity, we performed this assay by non-specifically adsorbing DLL4 proteins to 96-well tissue culture plates. This facile method requires minimal manipulation of proteins and is an established method for stimulating Notch activation *in vitro*. Plates were coated with various concentrations of WT DLL4 or Delta^MAX^ and Notch1 reporter cells were cultured on the ligand-coated surfaces. After 24 hours, fluorescence was monitored by flow cytometry. Fitting the data revealed that EC50 values for Notch1 activation by WT DLL4 and Delta^MAX^ were 1.7 μM and 6.8 nM, respectively, which corresponds to a 250-fold improvement in Notch activation efficiency (Fig. 3a). Moreover, the E_max_ induced by Delta^MAX^ was 20% greater than that induced by WT DLL4, indicating that Delta^MAX^ is capable of activating Notch signaling with greater amplitude and potency than WT DLL4.

**Fig 3.**
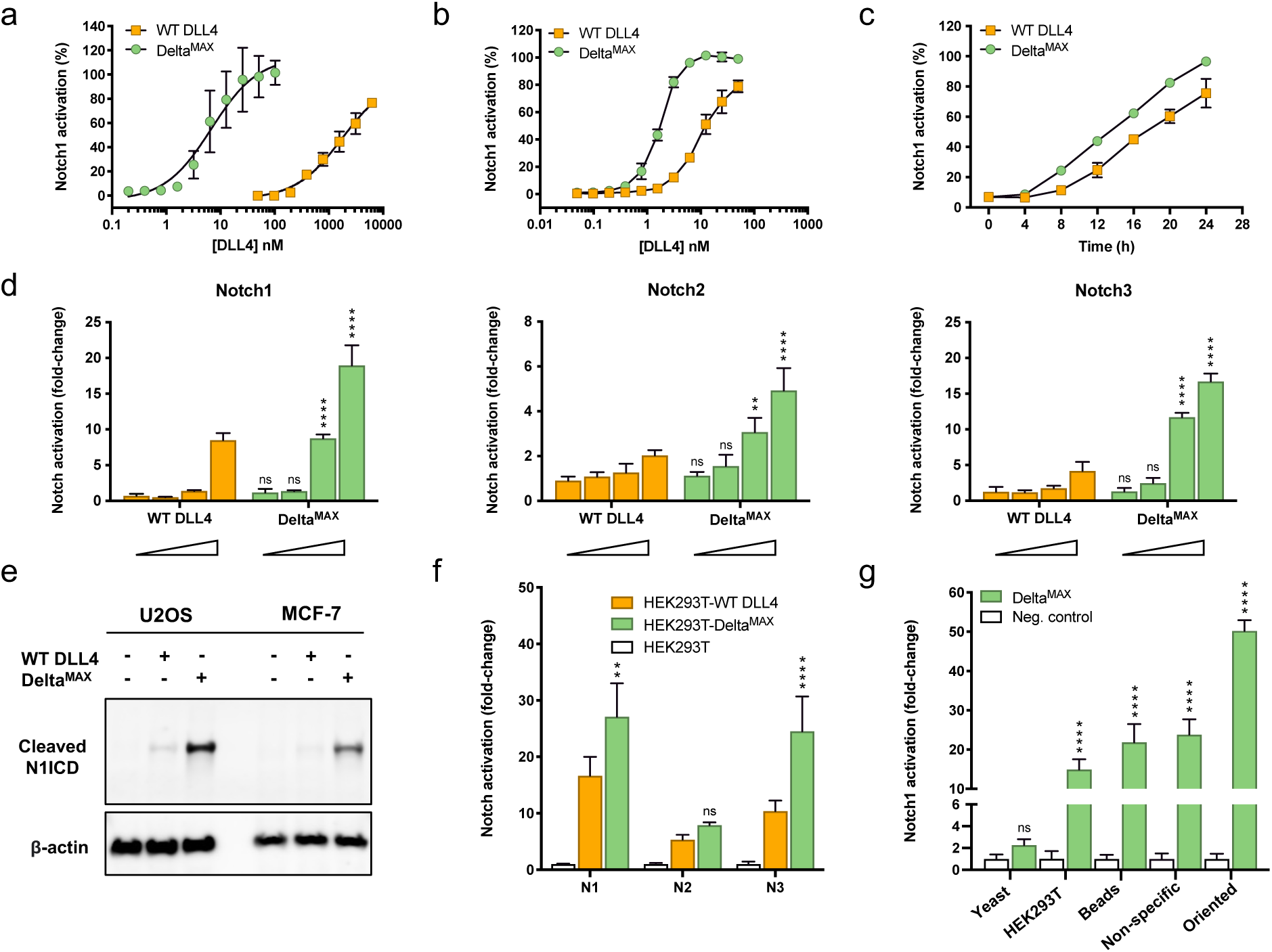
Delta^MAX^ is more potent Notch agonist than WT DLL4. (**A**) Dose-titration assay comparing the Notch1 reporter activity stimulated by WT DLL4 and Delta^MAX^ proteins that were non-specifically adsorbed to tissue culture plates. (**B**) Dose-titration assay comparing the Notch1 reporter activity stimulated by WT DLL4 and Delta^MAX^ proteins. DLL4 ligands were C-termini biotinylated and coupled to streptavidin-coated plates. (**C**) Time-course experiment comparing Notch1 reporter activity stimulated by WT DLL4 or Delta^MAX^ immobilized at 50 nM. (**D**) A luciferase reporter assay was used to measure Notch1, Notch2, or Notch3 activation by WT DLL4 and DLL4^MAX^. Protein concentrations were a 10-fold serial dilution from 50 nM to 0.05 nM. (**E**) Detection of endogenous cleaved Notch intracellular domain (N1ICD) in U2OS and MCF-7 cell lines by Western blot. Cells were Notch activated with 50 nM of WT DLL4 and Delta^MAX^ immobilized in streptavidin plates. (**F**) Co-culture assay comparing Notch1 activation by Delta^MAX^ and WT DLL4. Notch1 reporter cells and ligand-expressing cells were cultured at a 1:1 ratio. (**G**) Notch1 luciferase reporter cells were stimulated with Delta^MAX^ protein in different formats. Yeast expressing Delta^MAX^ were co-cultured in 10:1 ratio with reporter cells; HEK293T cells stably expressing Delta^MAX^ were co-cultured in 1:1 ratio; Delta^MAX^-coated streptavidin beads were mixed in 20:1 ratio; 100 nM Delta^MAX^ was non-specifically adsorbed to surfaces prior to addition of reporter cells; 10 nM C-terminally biotinylated Delta^MAX^ was immobilized on streptavidin-coated plates. Notch activation was normalized to the corresponding controls. Delta^MAX^ statistics are referred to WT DLL4. *P<0.05, **P<0.01, ****P<0.0001, ns: not significant (Two-way ANOVA).

We next tested whether C-terminal anchoring of DLL4 proteins improves signaling output. This strategy mimics the orientation the proteins adopt on the cell surface and was achieved by biotinylating the ligand C-termini prior to attachment to streptavidin-coated plates. For both WT DLL4 and Delta^MAX^, oriented coupling induced a higher maximum level of Notch1 activation than non-specific adsorption (Fig. 3b). In this format, Delta^MAX^ was also more potent than WT DLL4 and activated Notch1 with an EC50 of 1.7 nM compared to 16 nM for WT DLL4. We also performed a time-course experiment to evaluate the signaling kinetics of Delta^MAX^. We determined that Delta^MAX^ activates Notch1 more rapidly than WT DLL4, and that this effect is most pronounced in the first 4 to 8 hours following ligand stimulation (Fig. 3c). These findings indicate that the high-affinity Delta^MAX^ ligand potently activates Notch regardless of coupling strategy, and that oriented coupling maximizes the efficiency of Notch activation through DLL4 ligands.

### Delta^MAX^ strongly activates multiple human Notch receptors

We tested the ability of Delta^MAX^ to activate different human Notch receptor subtypes using an established luciferase reporter assay. For this assay, we used U2OS cells stably expressing chimeric Notch1, Notch2, or Notch3 proteins (Extended Data Fig. 6b)^32^. Notch4 was excluded from our study since it has been suggested that Notch4 inhibits signaling and it is not responsive to ligand-mediated activation *in vitro*^33^. To compare the signaling of each ligand, we cultured Notch1, Notch2, and Notch3 reporter cells with C-terminally anchored WT DLL4 or high-affinity Delta^MAX^ (Fig 3d). When we stimulated Notch1 cells with WT DLL4, we observed a dose-dependent increase in Notch signaling. The Delta^MAX^ variant more potently activated Notch1 across all concentrations and induced 2-fold higher levels of Notch1 signaling relative to WT DLL4 at the highest concentration tested (50 nM). Delta^MAX^ also activated Notch2 and Notch3 more potently than WT DLL4. For Notch2 and Notch3, relative Delta^MAX^ signaling was increased by 2.5-fold and 3-fold, respectively. These functional data indicate that enhanced affinity of Delta^MAX^ leads to increased signaling through Notch1-3, and that Delta^MAX^ is a viable tool for potently activating multiple human Notch receptors.

To determine whether Delta^MAX^ stimulates increased activation in cells expressing endogenous levels of Notch1, we compared the signaling of Delta^MAX^ and WT DLL4 in U2OS (osteosarcoma epithelial) cells and MCF-7 (breast cancer). U2OS and MCF-7 cells were each stimulated for 24 hours using C-terminally anchored WT DLL4 and Delta^MAX^ and Notch1 activation was detected by Western blotting for the cleaved Notch1 ICD (Fig. 3e). We found that Delta^MAX^ activated Notch signaling much more potently than WT DLL4 in both cell lines, indicating that Delta^MAX^ is a more potent ligand regardless of Notch expression level.

Thus far, all our signaling assays were performed using purified recombinant ligands. To evaluate the potency of Delta^MAX^ when expressed as a full-length transmembrane protein in cells, we performed a co-culture Notch signaling assay. We generated HEK293T cell lines stably expressing either full-length WT DLL4 or Delta^MAX^ (Extended Data Fig. 7b) and then co-cultured the ligand-expressing cells with Notch1, Notch2, or Notch3 reporter cells (Extended Data Fig. 6b) at a 1:1 ratio. In this co-culture format, we determined that the E_max_ of Delta^MAX^ signaling through Notch1 and Notch3 was ∼2-fold higher than WT DLL4. The luciferase signal for Notch2 was low for both ligands, although only Delta^MAX^ induced a significant level of Notch2 activation compared to unstimulated cells (*P* = 0.037) (Fig. 3f).

### Comparison of Delta^MAX^ signaling on plates, beads, and cells

A myriad of ligand-presentation platforms have been developed to activate mechanosensitive Notch receptors, including ligand-coated plates, ligand-coated beads, antibody-clustered ligands, ligand-infused matrices, and ligand-overexpressing stromal cell lines^13, 14, 18, 30, 34–36^. To our knowledge, no single experiment has directly compared the level of Notch signaling induced across different ligand-presentation formats. Here, we compared the signaling of Delta^MAX^ in five different formats to determine an optimal strategy for maximizing Notch activation. Fluorescent Notch1 reporter cells were cultured with yeast-displayed Delta^MAX^, 293T cells-expressing Delta^MAX^, Delta^MAX^-coated microbeads, plates coated adsorbed with Delta^MAX^ (non-specific coupling), and streptavidin plates coated with C-terminally biotinylated Delta^MAX^ (oriented coupling) (Fig. 3g, Extended Data Fig. 8). We determined that yeast-displayed Delta^MAX^ did not induce a significant change in Notch1 activation, which may be due to the inability of cell wall-tethered ligands to undergo endocytosis, or because the low mass of the yeast cells provides insufficient resistance. However, the remaining formats induced 15- to 50-fold increases in Notch1 activation relative to controls. The co-culture method induced the lowest level of Notch1 activation (15-fold), followed by Delta^MAX^-coated beads (21-fold), non-specific adsorption (23-fold), and oriented coupling (50-fold) (Fig. 3g). Taken together, our data reveal that recombinant protein-based formats (plate- or bead-bound ligands) induced higher levels signaling than cell-based methods. Furthermore, the increased E_max_ of C-terminally anchored Delta^MAX^ suggests that the greater accessibility of the ligand-binding C2-EGF3 domains is optimal for maximal Notch stimulation.

### Soluble Delta^MAX^ is a potent pan-Notch inhibitor

In the absence of membrane anchor, DLL4 ECDs are predicted to block ligand-mediated Notch activation by functioning as soluble decoys. However, WT DLL4 binds to human Notch receptors with micromolar affinities that fall outside of the expected range for effective pharmacological inhibition. We hypothesized that our high-affinity DLL4 variants would overcome the affinity-limitations of WT DLL4 to function as potent Notch antagonists. To test this hypothesis, we cultured fluorescent Notch1 reporter cells on C-terminally anchored WT DLL4-coated plates in the presence of soluble competitors. We added increasing concentrations of soluble WT DLL4, DLL4.v1, DLL4.v2, and Delta^MAX^ ECDs to the cells and then monitored reporter signal after 24 hours. We determined that, among the four proteins, only soluble Delta^MAX^ was able to efficiently block plate-bound WT DLL4-induced Notch1 activation and inhibited signaling with an IC50 of 0.6 nM. By contrast, administration of soluble WT DLL4 only reduced reporter activity by ∼25% at the highest concentration tested (300 nM). DLL4.v1 and DLL4.v2 were more potent inhibitors than WT DLL4, but less potent than Delta^MAX^, and neither DLL4.v1 or DLL4.v2 fully blocked Notch1 activation (Fig. 4a).

**Fig 4.**
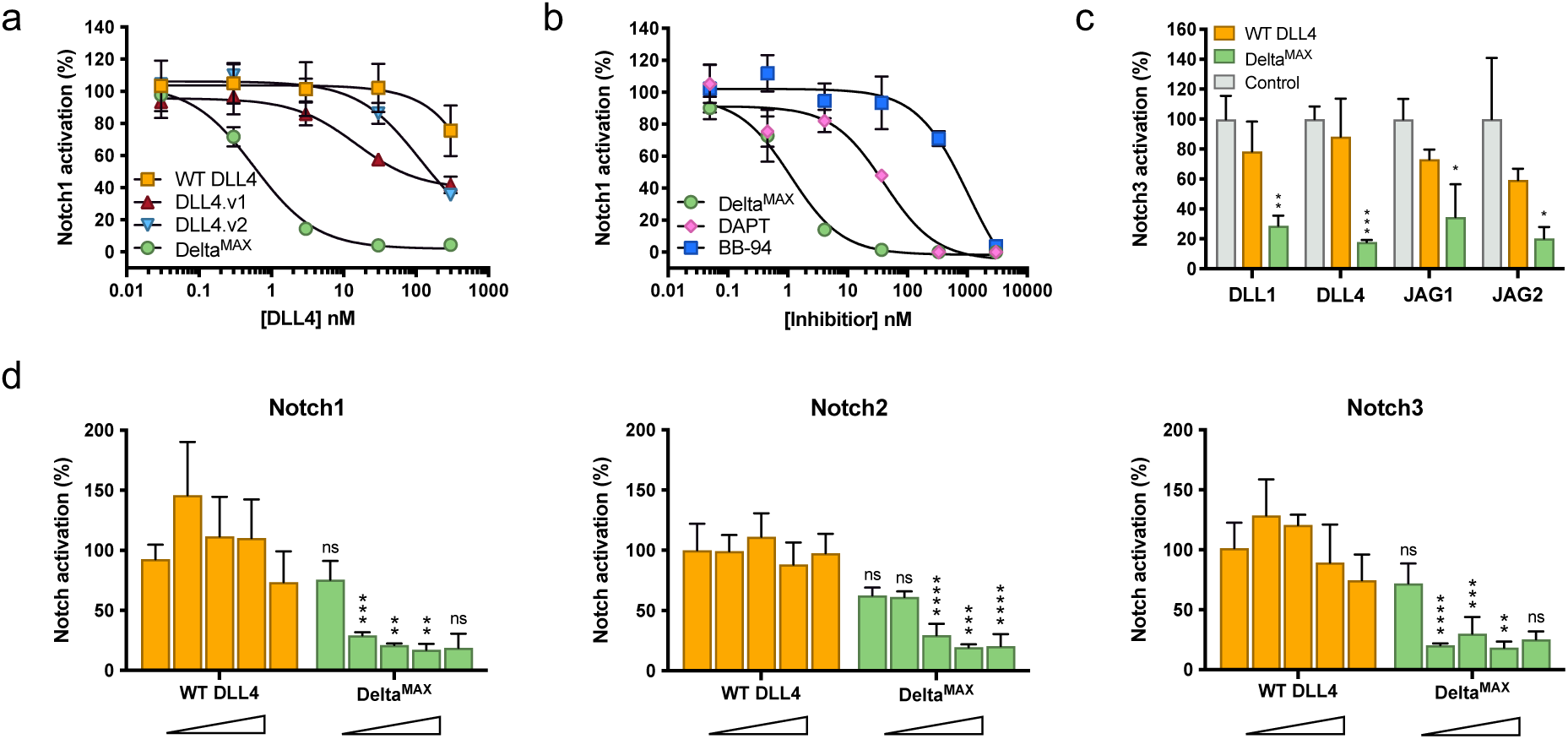
Delta^MAX^ is a pan-Notch inhibitor. (**A**) Dose-titration assay measuring the inhibition of Notch1 reporter activity by soluble DLL4 variants. Notch1 signaling was activated by culturing cells on streptavidin plates coated with 50 nM WT DLL4. (**B**) Dose-titration assay comparing the inhibition potency of DAPT, BB-94, and soluble Delta^MAX^. Notch1 signaling was activated by culturing cells on streptavidin plates coated with 50 nM WT DLL4. (**C**) Soluble WT DLL4 and Delta^MAX^ at 3 µM were tested for their ability to inhibit Notch1 activation by 293T cells overexpressing DLL4, DLL1, JAG1, or JAG2. (**D**) Dose-titration assay comparing the ability of WT DLL4 and Delta^MAX^ to inhibit activation of Notch1-3 by WT DLL4. 10-fold serial dilutions starting at 3,000 nM of soluble DLL4 or Delta^MAX^ were added to Notch1, Notch2, and Notch3 luciferase reporter cells. Notch signaling was activated by culturing cells on streptavidin plates coated with 50 nM WT DLL4. *P<0.05, **P<0.01, ***P<0.001, ****P<0.0001, ns: not significant (Two-way ANOVA).

Most small molecule Notch inhibitors are non-specific and target multifunctional ADAM10/17 or gamma-secretase proteases required for Notch activation. Therefore, we compared the inhibition potency of our Notch-selective Delta^MAX^ protein to the established gamma-secretase inhibitor DAPT and BB-94, a broad-spectrum metalloprotease inhibitor, BB-94 that inhibits ADAM10 (Fig. 4b). We determined that Delta^MAX^ inhibited Notch1 activation with an IC50 of 1 nM, which was 40-fold more potent than DAPT (IC50 of 40 nM) and 1000-fold more potent than BB-94 (IC50 of 1000 nM). We also compared the inhibition of Delta^MAX^ and DAPT in a co-culture assay. Both DAPT and Delta^MAX^ inhibited cellular DLL4 less potently than plate-bound DLL4, and the IC50 of DAPT and Delta^MAX^ were 100 and 400 nM, respectively (Extended Data Fig. 9). Thus, Delta^MAX^ functions as a potent and selective inhibitor of ligand-mediated Notch signaling, and the potency of inhibition depends on the format in which the wild-type ligand is presented.

We also tested the ability of Delta^MAX^ to inhibit signaling by ligands other than DLL4. First, we generated stable HEK293T cell lines expressing DLL1, DLL4, Jag1, and Jag2 and confirmed ligand-expression levels by antibody staining (Extended Data Fig. 10). We then co-cultured DLL1, DLL4, Jag1 or Jag2 cells with Notch3 luciferase reporter cells that were pre-mixed with a constant (3 μM) concentration of soluble WT DLL4 or Delta^MAX^ (Fig. 4c). We determined that Delta^MAX^ effectively inhibited productive signaling by DLL1, DLL4, Jag1, and Jag2, with a reduction in reporter signal ranging from 70-85%. On the other hand, WT DLL4 only weakly inhibited Notch activation through the four ligands, reducing reporter signal by 10-40%.

Lastly, we assessed whether Delta^MAX^ could inhibit signaling through different human Notch paralogs. We incubated Notch1, Notch2 and Notch3 luciferase reporter cell lines with various concentrations of soluble WT DLL4 or Delta^MAX^, and then cultured the cells on plates coated with DLL4-coated plates to monitor Notch activation (Fig. 4d). We determined that addition of the Delta^MAX^ competitor led to dose-dependent decreases in signaling for Notch1, Notch2, and Notch3, and that WT DLL4 did not significantly inhibit Notch activation at all concentrations tested.

### Delta^MAX^ increased human CD8^+^ T cell proliferation and effector markers

Notch signaling contributes to several aspects of immunobiology and influences the proliferation, differentiation, and antitumor function of CD8^+^ T cells^37–41^. Therefore, we developed an assay to test the effect of Delta^MAX^ on activated T cells (Fig. 5a). As stromal cells are an established platform for Notch stimulation in T cells, we generated artificial antigen presenting K562 cells (aAPCs) expressing the Fc receptor CD32 (renamed K32 cells), and similar surface levels of WT DLL4 or Delta^MAX^ (Fig. 5b)^42^. We then co-cultured these ligand-expressing cells with primary T cells in the presence of increasing concentrations of the T cell-activating antibody OKT3. CD32 expression enabled OKT3 Fc binding to K32 cells and subsequent CD3-TCR activation on T cells. Human CD8^+^ T cells were negatively isolated from peripheral blood and co-cultured with K32 cells pre-loaded with increasing concentrations of OKT3 to analyze T cell proliferation and IFNγ secretion (Fig. 5c, 5d). Delta^MAX^ treatment enhanced the percentage of proliferating CD8^+^ T cells between 27% and 15% along the OKT3 concentrations assayed (*P* = 0.032), while WT DLL4 did not show a significant improvement over the stimulation with K32 cells lacking DLL4 ligands (mock) (Fig. 5c). Additionally, Delta^MAX^ increased IFNγ secretion from CD8^+^ T cells 10-15% compared with WT DLL4 treatment (*P* = 0.047) (Fig. 5d).

**Fig 5.**
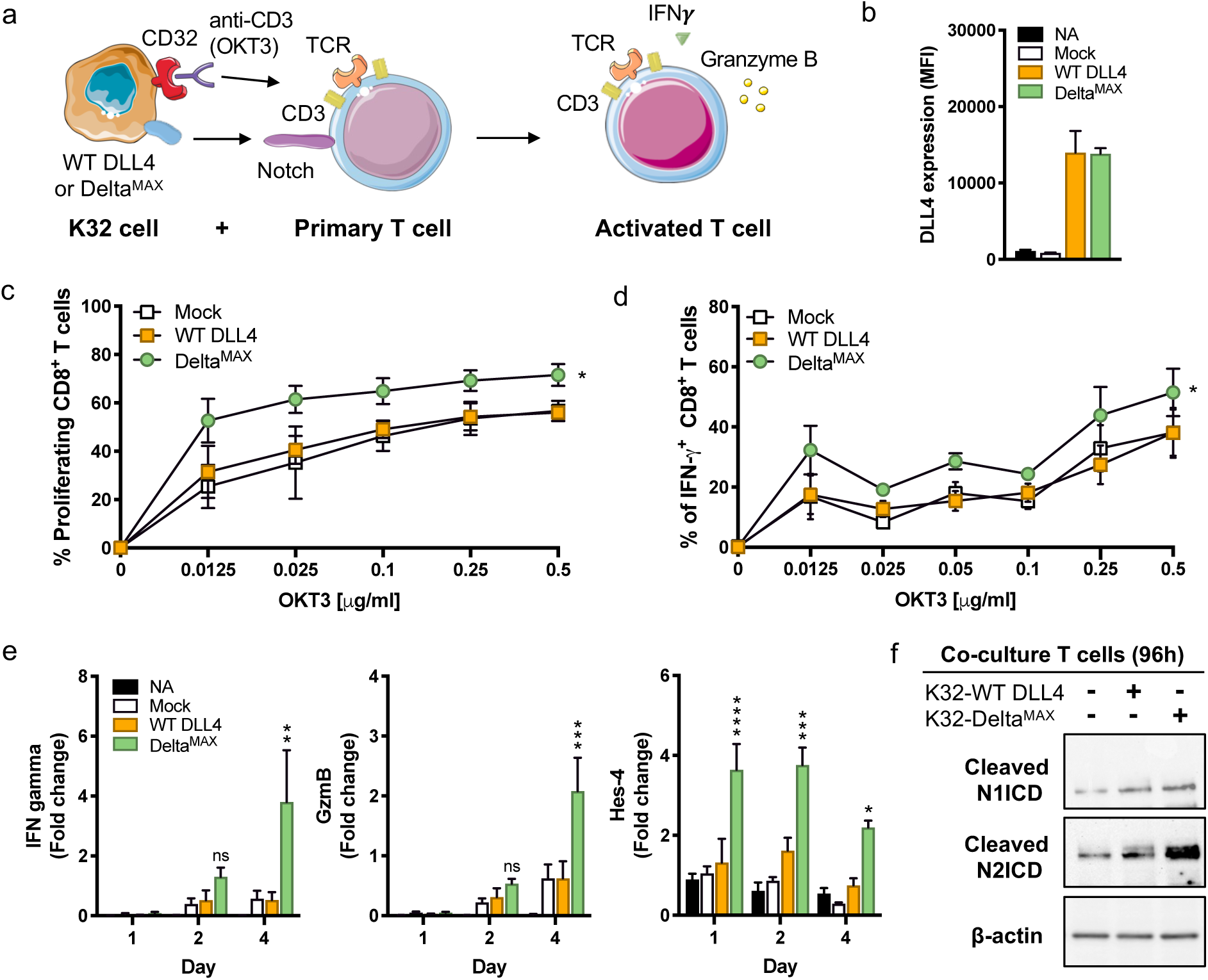
Translational strategy to transiently boost Notch signaling in human CD8^+^ T cells. (**A**) Human PBMCs CD8^+^ T cells were co-cultured with K32 artificial antigen-presenting cells (aAPCs) engineered to express WT DLL4 or Delta^MAX^ for simultaneous Notch and TCR stimulation. (**B**) aAPCs were assayed for expression of WT DLL4 or Delta^MAX^ using flow cytometry. K32 cells were loaded with OKT3 antibody using a gradient of concentrations starting at 0.0125 µg/ml, and then co-cultured with CD8^+^ T cells in ratio 10:1. After 96h, CD8^+^ T cells were analyzed for proliferation (**C**) and IFNγ secretion (**D**). **(E)** T cells were stimulated for 24, 48, and 96h to quantify mRNA levels of intracellular activation markers by real-time PCR, including IFNγ, Granzyme B and Hes-4. (**F**) K32 cells expressing WT DLL4 or Delta^MAX^ were co-culture with human CD8^+^ T cells (ratio 1:10) for detection of Notch activation by Western blot. Co-cultured CD8^+^ T cells were positively sorted by MACS column after cultivation with K32 cells, and nuclear extracts were harvested after 96 hours. Cleavage of Notch1 and Notch2 were examined by Western blot with specific antibodies. β-actin was used for loading control. Western blot images are representative results of 3 biological repeats. NA: non-activated T cells. Statistics for Delta^MAX^ are referred to WT DLL4. *P<0.05, **P<0.01, ***P<0.001, ***P<0.0001, ns: not significant difference (Two-way ANOVA). T cell proliferation and IFNγ secretion data were analyzed by unpaired t tests.

Next, we used RT-PCR to quantify the mRNA levels of Notch and T cell effector related markers over four days of CD8^+^ T cell stimulation with K32 loaded with 0.0125 μg/mL OKT3. We determined that Delta^MAX^-stimulated T cells had increased levels of effector mediators IFNγ and Granzyme B, as well as elevated expression of Notch target Hes-4 compared to counterparts conditioned with WT DLL4 or mock (Fig. 5e). Lastly, we performed a western blot analysis to compare Notch1 and Notch2 activation in T cells that were stimulated with WT DLL4 or Delta^MAX^. Using an antibody to the cleaved Notch1 or Notch2 ICD, we determined that activation of both Notch1 and Notch2 was increased for Delta^MAX^-treated cells relative to WT DLL4, and the difference in Notch2 activation was more prominent than that of Notch1 (Fig. 5f). Thus, results show the promotion of CD8^+^ T cell function induced by aAPCs expressing Delta^MAX^.

## DISCUSSION

Recombinant DLL and Jag ligand ECDs bind weakly to Notch receptors and are inefficient modulators of Notch signaling. We engineered the synthetic Delta^MAX^ ligand to be a versatile tool for activating or inhibiting mammalian Notch receptors, depending on the manner in which it is administered. Compared to WT DLL4, immobilized or cellular Delta^MAX^ exhibits varying degrees of increased signaling potency, and soluble Delta^MAX^ functioned as a Notch-specific inhibitor when administered as a soluble decoy. Furthermore, the increased thermostability (ΔT_m_ 11°C) and expression (2-fold higher yield) of Delta^MAX^ are both favorable properties for protein biologics.

Our functional studies revealed that Delta^MAX^ activates Notch more potently than WT DLL4 in plate-bound, bead-bound, and cellular formats. Among the methods tested, we determined that C-terminally anchored Delta^MAX^ induced the highest levels of Notch signaling, followed by non-specifically adsorbed ligands, ligand-coated beads, and cellular ligands. The largest difference in signaling potency between WT DLL4 and Delta^MAX^ was observed when ligands were non-specifically adsorbed to plastic. The WT DLL4 protein was a poor Notch activator in this format (EC50 1.7 μM), while Delta^MAX^ was remarkably potent (EC50 6.8 nM). The strong signaling activity can enable entire flasks or tissue culture plates to be pre-conditioned with Delta^MAX^ as a high-throughput means of activating Notch in cultured cells. Furthermore, non-specific adsorption requires low quantities of protein and can be performed with unmodified ligand ECDs, making this method highly accessible for a wide range of biotechnological applications.

Soluble DLL4 ECDs do not activate Notch because they are unable to exert the requisite pulling force. Instead, these ECD are predicted to inhibit signaling by competing for the ligand-binding site on the Notch receptor. We demonstrated that, even at high-concentrations, the micromolar binding affinity of WT DLL4 prevents it from functioning as an effective Notch antagonist. On the other hand, Delta^MAX^ potently inhibited Notch activation by all four activating ligands (DLL1, DLL4, Jag1, and Jag2). The broad-reactivity profile of Delta^MAX^ also enabled it to inhibit ligand-mediated activation of Notch1, Notch2, and Notch3. Current strategies for pan-Notch inhibition require the use of gamma-secretase or ADAM10/17 inhibitors, which block the cleavage of numerous cellular substrates in addition to Notch. Alternatively, antibodies targeting the NRR domain have been used to block signaling by individual Notch receptor subtypes. We found that Delta^MAX^ inhibited Notch signaling with similar potency to the established gamma-secretase inhibitor DAPT. Therefore, this ability of Delta^MAX^ to block signaling by all Notch receptors makes it a unique pharmacological tool as a highly selective, pan-Notch inhibitor.

Notch signaling is central regulator of T cell biology and influences T cell activation, proliferation, and cytolytic effector functions^37–40, 43, 44^. We previously demonstrated that constitutive Notch activation by overexpression of NICD enhances the antitumor activity of adoptively transferred murine CD8^+^ T cells^45^. As this method required genetic modification of the T cells, we aimed to develop an *ex vivo* platform for transient Notch activation using Delta^MAX^. In the present study, we determined that co-culturing activated T cells with K32 cells expressing Delta^MAX^ induces increased T cell proliferation and expression of effector mediators compared to co-culture with WT DLL4. This approach could be potentially beneficial for strategies of expansion of anti-tumor CAR-T or TILS for further adoptive transfer by triggering Notch-related metabolic and anti-tumor signals.

Various ligand presentation strategies have been developed for activating Notch in T cells, and in these systems, the dosage of Notch signaling has proven to be important for achieving optimal outcomes. For example, low doses of DLL1 stimulate the differentiation of hematopoietic stem and progenitor cells (HSPCs) into mixed populations of lymphoid and myeloid precursors, whereas high-doses promote the differentiation of only lymphoid precursors^46^. Additionally, stromal cells that express DLL4, which binds to Notch1 with higher affinity than DLL1^13^, more efficiently drives T cell lymphopoiesis^35^. Given the tunable outcomes of T cell behavior, we anticipate that the ultra-potent Delta^MAX^ ligand will be a powerful tool for achieving ideal levels of Notch activation for T cell-related applications.

## ACKNOWLEDGMENTS

We would like to thank Dr. Eric Lau, Dr. Gregory Watson and Daniel Lester for their helpful assistance in generating DLL4 stable cell lines. We also thank Dr. Timothy Tran from Moffitt Chemical Biology Core for helping with DSF studies, and Dr. Qianqian Ming for helpful suggestions. D.G. and V.L. were supported by R35GM133482 and an award from the Rita Allen Foundation. D.A. was supported by the Sigrid Juselius Foundation. S.C.B. is supported by R35CA220340. P.C.R. and S.D. were supported by R01CA184185, R01CA233512; R01CA262121; P01CA250984 Project #4; and P30CA076292; and Florida Department of Health grant #20B04. Support for shared resources was provided by the Moffitt Cancer Center Support Grant NIH P30CA076292.

## AUTHOR CONTRIBUTION

V.C.L. and D.G.P. wrote the manuscript. V.C.L., D.G.P, D.A., and P.C.R. designed the experiments. D.G.P. performed the protein purifications, yeast display selections, binding studies, and signaling assays. S.D. performed and designed the T cell experiments with assistance from P.C.R. E.M. contributed to library design and cloning. D.A. performed the detection of Notch activation assays on U2OS and MCF-7 cells by Western blot. E.D.E. and S.C.B. provided U2OS reporter cells and assisted with luciferase experimental procedures. S.C.B., P.C.R., and V.C.L. edited and reviewed the manuscript. V.C.L. supervised and conceived the project.

## MATERIAL AND METHODS

### Reagents, media, and cell lines

Mammalian cells were cultured at 37°C, with a humidified atmosphere of 5% CO_2_, washed with Dulbecco’s Phosphate Saline Buffer (DPBS, Corning), and detached with Trypsin-EDTA 0.25% (Gibco) for cell sub-culturing or cell-based assays. Notch reporter cell lines CHO-K1 hNotch1-Gal4 were a gift from Dr. Michael Elowitz (California Institute of Technology)^1^. HEK293T, U2OS and MCF-7 cells were cultured in high-glucose Dulbecco Eagle’s Minimal Essential Medium (DMEM, Cytiva) supplemented with 10% Fetal Bovine Serum (FBS, peak serum) and 2% penicillin/streptomycin (Gibco). The MCF-7 breast cancer cell line was a gift from Dr. Eric Lau (Moffitt Cancer Center). Hygromycin 100 µg/ml was added to U2OS reporter^2^ cell cultures to maintain homogeneous populations of Notch-expressing cells. CHO-K1 hNotch1-Gal4 cells were cultured in Minimum Essential Medium Eagle-alpha modification (α-MEM, Cytiva) supplemented with 10% Fetal Bovine Serum (FBS, peak serum), 2% penicillin/streptomycin (Gibco), 400 µg/mL of Zeocin (Alfa aesar) and 600 µg/mL of Geneticin (Gibco). K32 cells (K562 cell line expressing CD32) were a gift from Dr. Jose Conejo-Garcia (Moffitt Cancer Center), and human CD8^+^ T cells were PBMCs (collected from LifeSouth Blood Bank, Gainesville, FL). Cells used for T cell experiments were cultured at 37°C, and 5% CO_2_, in RPMI-1640 (Gibco) supplemented with 2 mM L-glutamine, 10 mM HEPES, 150 U/mL streptomycin, 200 U/mL Penicillin, 20 mM β-mercaptoethanol and 10% heat-inactivated FBS (GeminiBio). All cell lines were tested negative for mycoplasma (ATCC), and stable cell lines were validated by sequencing (Genewiz) after genomic DNA extraction (Invitrogen). Antibodies used in this study were: anti-hNotch1 PE (Biolegend), anti-hNotch2 AF647 (Biolegend), anti-hNotch3 PE (Biolegend), anti-hDLL4 PE (Biolegend), anti-hDLL1 PE (Biolegend), anti-hJagged1 AF647 (Biolegend), anti-hJagged2 PE (Biolegend), and anti-HA tag AF488 (Cell Signaling Technologies), anti-hCD8a Bv785 (Biolegend), anti-hIFNγ APC (Biolegend), goat anti-mouse IgG (secondary antibody, KPL), goat anti-rabbit IgG (secondary antibody, Vector laboratories), anti-cleaved Notch1 (Val1744, Cell Signaling Technology), anti-Notch2 (D76A6, Cell Signaling Technologies), anti-β-Actin (Cell Signaling Technology and Sigma).

### Protein expression and purification

The minimum binding region of Delta-Like 4 (DLL4) fragments (N-EGF5) were cloned into the pAcGp67A vector containing an N-terminal gp67 signal peptide and C-terminal 8xHis-tag. Non-biotinylated proteins were human wild-type DLL4 (N-EGF5, residues 27-400), DLL4.v1 (N-EGF5), DLL4.v2 (N-EGF5), and DLL4^MAX^ (N-EGF5). Notch biotinylated fragments were cloned into pAcGp67A with a N-terminal gp67 signal peptide, a C-terminal 3C protease site followed by biotin-acceptor peptide tag (BAP-tag: GLNDIFEAQKIEWHE), and 6xHis-tag. In contrast, DLL4 biotinylated proteins contained BAP and His-tags, but lacked 3C site. Biotinylated proteins were human Notch1 (EGF8-12, residues 295-488), hNotch1 (EGF6-13, residues 237-549), hNotch2 (EGF6-13, residues 221-530), hNotch3 (EGF5-12, residues 197-505), hNotch4 (EGF7-12, residues 276-511), mouse Notch1 (EGF6-13, residues 221-526), WT DLL4 (N-EGF5, residues 27-400), and DLL4^MAX^ (N-EGF5, residues 27-400). All baculovirus-based constructs in this work were expressed for 48 hours by infecting HiFive *Trichoplusia ni* insect cell cultures (Expression Systems) at a density of 2×10^6^ cells/mL with infective baculovirus particles. Culture supernatants were harvested, and proteins purified by IMAC. Ni-NTA agarose resin (Qiagen) was washed with HEPES Buffered Saline (HBS: 20 mM HEPES pH 7.4, 150 mM sodium chloride; Buffer A) supplemented with 25 mM imidazole (plus 1 mM calcium chloride for Notch preps; Buffer A+C) and eluted with Buffer A (or Buffer A+C for Notch samples) containing 250 mM imidazole. Polishing was performed using a Superdex 200 Increase 10/300 GL column (GE) equilibrated in Buffer A for DLL4 proteins, or Superdex 75 Increase 10/300 GL column (GE) equilibrated in Buffer A+C for Notch proteins. Biotinylated proteins were site-specifically modified *in vitro* at their C-terminal BAP-tag, using BirA ligase (in-house prepared) and polished as described above. Protein purity and integrity were analyzed by SDS-PAGE using TGX 12% Pre-cast gels (Bio-Rad). All proteins were flash frozen in liquid nitrogen and stored at -80°C for later use.

### Affinity maturation of human DLL4 by directed evolution

#### Starting point of evolution and structure-guided library creation

Wild-type human Delta-like 4 (N-EGF3, residues 27-322) was cloned into a modified version of pCT302 an N-terminal fusion to an HA-epitope (YPYDVPDYA), c-Myc epitope (EQKLISEEDL), and Aga2. A third binding site for DLL4:Notch interaction was designed on the basis of the binding interface found in the crystal structure of rJagged1(N-EGF3):rNotch1(EGF8-12) complex^3^. The complex was analyzed with different computational tools (PDBePISA (EMBL-EBI), Consurf server (Biosof), and PyMOL (Schrödinger)). Nine interface residues were selected to create a structure-guided directed evolution library. Jagged1 homolog residues were identified in DLL4 and targeted for mutagenesis. The mutant library was generated using assembly PCR with the ten overlapping oligonucleotides below:

Lib1F: CAGCCTATCTGTCTTTCGGGCTGTCATGAACAGAATGGCTACTGCAGCAAGC
Lib2R: GCCAGCCTGGGCGGCAGAGGCACTCGGCCGGCTTGCTGCAGTAGCCATTC
Lib3F: TCTGCCGCCCAGGCTGGCAGGGCCGGCTGTGTAACGAATGCATCCCC
Lib4R: CCAGGGAGTGCTGCAGGTGCCGTGGCGACAGCC**<u>ADDAWN</u>**GGGGATGCATTCGTTAC ACAG
Lib5F: CACCTGCAGCACTCCCTGGCAATGT**MHT**TGTGATGAGGGCTGGGGAG,
Lib6R: GCAGTAGTTGAGATCTTGGTCACA**<u>AWRCWK</u>**GCCTCCCCAGCCCTCATCACA
Lib7F: GTGACCAAGATCTCAACTACTGC**<u>RVT</u>**CACCACTCCCCATGCAAGAATGGGGCAACG TGC
Lib8R: GGCGACAGGTGCAGGTGTAGCTTCG**<u>CNG</u>**CCCACT**<u>WTBAYK</u>**GCACGTTGCCCCATTC TTG
Lib9F: CTACACCTGCACCTGTCGCCCGGGCTACACTGGTGTGGACTGTGAGGGATCCTACCC ATAC
Lib10R: AGATAAGCTTTTGTTCTCCACCAGCGTAATCTGGAACATCGTATGGGTAGGATCCCT CAC.

The primers contained specific degenerate codons (bold underlined) and 20bp overhangs (Eurofins) to promote annealing in the PCR reaction. External primers were added to introduce a 40-50 bp recombination area between the mutant library region (EGF1-EGF3) and DLL4-pYAL linearized vector. The mutant library was *in vivo* reassembled into yeast by electroporation (BTX electroporator) with 10 µg of linearized DLL4-pYAL vector and 50 µg of the mutagenic library (ratio 1:5 vector:library). Yeast transformants were recovered in SDCAA and DLL4 surface displayed on *S. cerevisiae* EBY100 yeast as reported previously^3, 4^. The theoretical library diversity was 4x10E6, and we obtained an estimated library size of 8x10E6 variants (2-fold theoretical diversity).

### Yeast display selections

Before starting each selection round, induced yeast cultures were pelleted in logarithmic phase, stained with anti-HA antibody conjugated to AF488 (Cell Signaling Technologies), and expression of DLL4 on yeast surface analyzed by flow cytometry (BD Accuri C6 Plus). Selection rounds were performed in two phases: (1) a negative selection to remove non-specific binders targeting streptavidin and/or Alexa Fluor 647, and (2) a positive selection to rescue high-affinity variants specific for Notch. Negative selection was performed by combining 100 nM of streptavidin conjugated to Alexa Fluor 647 (SA647) and 250 µL anti-Alexa Fluor 647 magnetic beads (Miltenyi) on ice for 10 min. Next, the yeast library was washed once in selection buffer (HBS: 20 mM HEPES pH 7.4, 150 mM sodium chloride, 1 mM calcium chloride, 0.1% Bovine Serum Albumin (BSA) and 10 mM maltose), and magnetic beads SA647-coated were added. The tube was wrapped in foil and incubated at 4°C for 30 min. Following this step, the yeast was washed twice with selection buffer, resuspended, and flowed over a Magnetic-Activated cell sorting (MACS) LS separation column (Miltenyi) pre-equilibrated in selection buffer. The flowthrough was rescued, and positive selections were performed. For round zero (Naïve library), 8x10E7 cells (10x library diversity) were used for selection against biotinylated Notch1 (EGF8-12) and Notch3 (EGF5-13), independently. Tetramers were pre-made on ice for 10 min by mixing 450 nM of biotinylated Notch1 (EGF8-12) or Notch3 (EGF5-12) with 100 nM of SA-647 (ratio 4.5:1). Next, 100 nM tetramers were added to 500 µL of negatively selected yeast in selection buffer and incubated for 2 hours at 4°C. Yeast library stained with Notch tetramers was washed twice with 500 µL of selection buffer, and further incubated with 50 µL of anti-AF647 magnetic beads for 30 min at 4°C. After incubation, the yeast was washed twice with 500 µL of selection buffer and high-affinity binders isolated by MACS columns. The isolated yeast cells were pelleted, recovered in 3 mL of SDCAA for 48 hours and induced in SGCAA for next round. For round 1, the same conditions were repeated like in round zero using Notch1 (EGF8-12) and Notch3 (EGF5-12) 100 nM tetramers for positive selections. Isolated populations were rescued in 3 mL SDCAA after MACS sorting and induced in SGCAA. For round 2, 5x10E7 yeast cells were resuspended in 500 µL of selection buffer in the presence of 2 µM monomer of Notch1 (EGF8-12) and Notch3 (EGF5-12), independently. Tubes were wrapped in foil and incubated for 2 hours at 4°C. Yeast samples binding Notch were washed twice with 500 µL of selection buffer, and incubated in 500 µL of selection buffer supplemented with 100 nM of SA-647 for 30 min at 4°C. DLL4 high-affinity binders were isolated by MACS following the same approach described for rounds zero and 1. High-affinity binders for Notch1 and Notch3 were rescued in 3 mL of SDCAA for 48 hours and induced in SGCAA. For round 3, high-affinity populations were isolated by Fluorescent-Activated Cell Sorting (FACS, Sony Sorter SH800S). Only Notch3 (EGF5-12) was used since yeast-staining tests showed better population enrichment compared with Notch1 (EGF8-12). 1x10E8 yeast cells were resuspended into 500 µL of selection buffer with pre-made 100 nM tetramers of Notch3 (5-12) and anti-HA488 antibody (double staining). The tube was wrapped in foil and incubated for 2 hours at 4°C. Stained yeast was washed two times with 500 µL of selection buffer and analyzed by FACS. From the double-positive population (FIT-C+, APC+) showing expression of DLL4 (anti-HA488, FIT-C signal) and binding to Notch (SA-647, APC signal), a sorting gate was set at 1.8% of the maximum signal detected for APC channel and Notch binders isolated (Extended Data Fig 1b). The sorted yeast population was washed two times with 5 mL of selection buffer, pelleted, and recovered in SDCAA for 48 hours. Yeast cells were induced again in SGCAA for the last round of selection. For round 4, 5x10E7 cells were incubated with 0.2 μM monomers of Notch3 (5-12) for 2 hours at 4°C. Yeast cells were then washed twice with selection buffer, fluorescent-labeled with SA-647 and high-affinity populations isolated by MACS following the protocol described for previous rounds.

The isolated yeast populations obtained after round 4 were subjected to colony screen and plasmid recovery by using Zymoprep kit (Zymo Research). The plasmids were transformed into *E. coli* and 30 individual colonies sequenced with pYAL F (5’-AAATGATAACCATCTCGC) and pYAL R (5’-GGGATTTGCTCGCATATAGTTG) primers (Eurofins). Sequencing data analysis was performed using SnapGene software (Insightful Science).

### Generation of DLL4 variants

We first generated a variant named DLL4.v1 by engrafting five affinity enhancing mutations into human DLL4 scaffold that we previously reported for rat DLL4 affinity maturation (Site 1 + Site 2; G28S, F107L, I143F, H194Y, and L206P)^4^. Next, the mutations found in this work to create a third binding site on human DLL4 (Site 3; N257P, T271L, F280Y, S301R, Q305P) were combined with Site 1 + Site 2 mutations into one single DLL4 scaffold, creating an ultrapotent DLL4 protein, named Delta^MAX^ (Site 1 + Site2; G28S, F107L, I143F, H194Y and L206P, and Site 3; N257P, T271L, F280Y, S301R, Q305P).

### Thermal denaturation experiments by differential scanning fluorometry (DSF)

WT DLL4 and Delta^MAX^ (N-EGF5) were diluted at 5 µM in 20 µL of HBS buffer, and then combined with 10 μM (5x) SYPRO Orange (ThermoFisher Scientific) prepared in 100% DMSO (final DMSO concentration was 0.1%). This mix was equilibrated at room temperature for 30 min. Proteins were analyzed in a 96-well microtiter plate using a StepOnePlus RT-PCR system (Applied Biosystems) applying a continuous heating gradient from 25 to 99°C with a step of 1% of temperature/min. Data was normalized and represented as the mean of three independent technical replicates. Similar Tm values were obtained for the replica measurements.

### Determination of equilibrium dissociation constants (K_D_) by Surface Plasmon Resonance (SPR)

Equilibrium dissociation constants were determined by surface plasmon resonance using a Biacore T200 instrument (GE Healthcare). Approximately 100 resonance units (RU) of biotinylated Notch extracellular fragments (Notch1 (EGF6-13), Notch2 (EGF6-13), Notch3 (EGF5-12), Notch4 (7-12), and mouse Notch1(EGF6-13)) were immobilized on individual flow cells at 5 μl/min using a streptavidin-coated sensor chip (Series S Sensor chip SA, GE Healthcare). Three-fold serial dilutions of recombinant DLL4 proteins (N-EGF5) starting at 50 μM (WT DLL4), 18 μM (DLL4.v2), and 6 μM (DLL4.v1 and Delta^MAX^) were flowed over the chip at 25°C in SPR buffer (20 mM HEPES pH 7.4, 150 mM NaCl, 1 mM calcium chloride, 0.1% BSA supplemented with 0.005% surfactant P20). Association (“on-rate”) and dissociation (“off-rate”) phases were performed at 10 μl/min for 120 seconds and 300 seconds, respectively. The sensor chip was regenerated after each injection with 30-second washes of 2.25M Magnesium Chloride containing 25% Ethylene Glycol at 50 μL/min. Data collection rate was performed at 10Hz, and curves were reference-subtracted from a flow cell containing 100 RU of a negative control non-related protein (biotinylated mouse RNF43 PA domain). The maximum RU for each experiment was normalized to a value of 100% and plotted as a function of concentration using Prism 9 software (GraphPad). Steady-state binding and kinetic curves were fitted using the Biacore T200 evaluation software to a 1:1 Langmuir model to determine the K_D_ values.

### Generation of stable cell lines

Ectodomains of WT DLL4 and Delta^MAX^ (N-EGF8, residues 27-518) were cloned into an intermediary vector containing WT DLL4 signal peptide, juxtamembrane, transmembrane, and intracellular domain, fused to C-terminal Myc (EQKLISEEDL) and Flag (DYKDDDDK) tags. This vector was generated after cloning of a synthetic DNA fragment (Eurofins) into pLenti-C-Myc-DDK-IRES-Puro vector (Invitrogen). For stable cell lines expressing DLL1 (residues 1-723), Jagged1 (residues 1-1218), and Jagged2 (residues 1-1238), their CDS was cloned into pLenti-C-Myc-DDK-IRES-Puro vector fused to c-Myc and Flag C-termini tags. HEK-293T mammalian cells were used for lentiviral particle production and transductions (kindly donated by Dr. Eric Lau, Moffitt Cancer Center). Briefly, transfections were carried out with packaging vectors VSV-G and d8.9 (kindly donated by Dr. Eric Lau, Moffitt Cancer Center) in the presence of Polyethyleneimine (PEI) at ratio 4:1 (DNA:PEI). Lentiviral particles were harvested after 48-72 hours and used to transduce HEK293T cells using 1 μg/mL polybrene (Millipore). Antibiotic selection was performed using Puromycin at 5 μg/mL (Gibco) for 10-15 days. Puromycin resistant cells expressing WT DLL4 and Delta^MAX^ were detached with trypsin-free dissociation buffer (Gibco), stained in DPBS + 0.5% BSA with anti-hDLL4 PE antibody for 1 hour at 4°C, and then sorted by FACS (Sony sorter SH800S). Sorted cells (mean fluorescence intensity around 1x10E5- 3x10E5) were washed with DPBS and recovered in DMEM+10% FBS at 37°C and 5% CO_2_ for 2 weeks. Expression of WT DLL4 and Delta^MAX^ on HEK293T surface was confirmed by flow cytometry (BD Accuri C6 plus) staining the cell lines with anti-hDLL4 PE in DPBS + 0.5% BSA (Extended Data Fig 7b). DLL1, Jagged1, and Jagged2 cell lines were stained with specific antibodies (anti-hDLL1 PE, anti-hJagged1 AF647, and anti-hJagged2 PE) for 1 hour at 4°C and analyzed by flow cytometry (BD Accuri C6 plus); however, these cell lines did not require further sorting steps (Extended Data Fig. 10).

K32 cells (K562 cells expressing human CD32) were transduced with lentivirus expressing WT DLL4 or Delta^MAX^ by spin-infection using 8 μg/mL polybrene (Millipore). Seventy-two hours after transduction, cells were analyzed for DLL4 expression and further cultured in RPMI + 10% FBS containing 1 µg/ml of puromycin (Gibco) at 37 ⁰C and 5% CO_2_. DLL4 expression was monitored by flow cytometry (CytoFLEX II, Beckman Coulter) using anti-hDLL4 PE (Biolegend) until the culture reached ∼100% of positive population on PE signal.

### Notch activation assays with ligands non-specifically adsorbed to plates

On day one, 2-fold serial dilutions of non-biotinylated DLL4 recombinant proteins (N-EGF5) were prepared in 100 μL of DPBS and used to coat 96-well plates (Coastar) for 1 hour at 37°C. WT DLL4 started at 50 μM and Delta^MAX^ at 204 nM. Then, the wells were washed three times with 200 μL of DPBS to remove unbound proteins. Next, CHO-K1 Notch1-Gal4 cells in culture were detached with trypsin-EDTA 0.25% (Gibco) and manually counted. Appropriate dilutions were prepared in α-MEM media to ensure 30,000 CHO-K1 cells per well in 200 μL. Ligand-coated plates and reporter cell lines were incubated for 24 hours at 37°C and 5% CO_2_. On day two, CHO-K1 Notch1-Gal4 cells were washed with 200 µL DPBS, detached with 30 µL of trypsin-EDTA 0.25%, and quenched with 200 µL of α-MEM media. Finally, cells were resuspended and H2B-mCitrine signal was measured by flow cytometry (BD Accuri C6 plus). CHO-K1 Notch1-Gal4 and ligand-lacking wells were used as a negative control. The measurements represent the mean fluorescent intensity as percentage of Notch activation ± S.D. of three biological replicates. Notch activation was reference-subtracted from a well coated with 0.5% BSA containing CHO-K1 cells. Gating strategies for this experiment are detailed in Extended Data Fig. 7a.

### Notch activation with ligand-oriented coupling on streptavidin plates

For CHO K1 reporters, on day one, tissue culture 96-well plates (Coastar) were pre-treated with 100 µL of streptavidin at 10 µg/mL (VWR) for 1 hour at 37°C, washed three times with 200 µL DPBS (Corning), and blocked with 100 µL BSA 2% (VWR) for 1 hour at 37°C. Excess BSA was removed by washing three times with 200 µL DPBS (Corning), and 100 µL of DPBS containing biotinylated DLL4 proteins were coupled to streptavidin for 1 hour at 37°C (2-fold serial dilutions starting at 50 nM). Non-coupled DLL4 proteins were removed by washing three times with 200 µL DPBS. Next, Notch reporter cells in culture were detached with trypsin-EDTA 0.25% (Gibco), and 30,000 cells in α-MEM media (200 µL) were added into individual wells. Assay plates were incubated for 24 hours at 37 °C, and 5% CO_2_. On day two, CHO-K1 Notch1-Gal4 cells were washed with 200 µL DPBS, detached with 30 µL of trypsin-EDTA 0.25%, and 200 µL of α-MEM media added. Cells were resuspended and Notch activation was measured as a function of fluorescence by flow cytometry (BD Accuri C6 plus). CHO-K1 Notch1-Gal4 cells and lacking-ligand wells were used as a negative control. The measurements represent the mean fluorescent intensity as percentage of Notch activation ± S.D. of three biological replicates. Notch activation was reference-subtracted from a well coated with a non-related control protein (PA domain of RNF128).

For U2OS reporter cells, approximately 5x10E5 cells were plated into 6-well plates (Falcon) and let to adhere overnight at 37 °C, 5% CO_2_ in DMEM media (Cytiva). On day two, media was replaced with serum-free DMEM and cells transfected with luciferase vectors. Cationic lipoplexes were generated for 15 min at room temperature by mixing 10 µL of Lipofectamine 2000 (Invitrogen), 2.5µg of Gal4-firefly luciferase reporter, and 50 ng of pRL-TK *Renilla* luciferase vector as internal control (ratio 1:5, Lipo:DNA). On day three, streptavidin plates were coupled with DLL4 ligands like described above using 10-fold serial dilutions starting at 50 nM, and transfected U2OS cells were detached with Trypsin-EDTA, washed with DPBS, and counted in the microscope. Twenty-thousand reporter cells were added to wells using 200 µL of DMEM media containing 2 µg/mL of doxycycline (Sigma) to induce Notch-Gal4 expression. Next, signaling plates were incubated for 24 hours at 37 °C and 5% CO_2_. On day four, luciferase firefly and *Renilla* signals were determined with Dual-Glo luciferase assay system (Promega) using a GloMax luminometer (Promega). A luminescence ratio firefly:*Renilla* was determined for every well and normalized to the signal of a non-related control protein (PA domain of RNF128). Notch activation is represented as the mean fold-change over the control ± S.D. of three independent biological replicates.

### Kinetics for Notch activation assay

96-well tissue culture plates (Coastar) were pre-coated using ligand-oriented coupling as described above with 50 nM of DLL4 biotinylated variants. CHO-K1 reporter cells were detached with trypsin-EDTA, washed with DPBS, and 200 µL of α-MEM media containing 30,000 cells were added into wells. Notch activation was analyzed as end-points for 0, 4, 8, 12, 16, 20, and 24-hour measurements. Desired time point wells containing CHO-K1 reporter cells were washed once with 200 µL DPBS, detached with trypsin-EDTA, and quenched with 200 µL of α-MEM media. Next, reporter cells were resuspended and analyzed by flow cytometry reading the fluorescence of H2B-mCitrine (BD Accuri C6 Plus). Every time point represents the mean fluorescence intensity of Notch activation as percentage ± S.D. of three independent biological replicates. Notch activation was reference-subtracted from a well containing streptavidin, BSA and CHO-K1 cells.

### Notch activation in co-culture format with HEK293T

On day one, U2OS reporter cells (signal receiver cells) were detached with trypsin-EDTA, washed with DPBS, and 5x10E5 cells were added to 6-well plates (Falcon) for transfection with luciferase reporter vectors. On day two, U2OS cells were transfected as described above. On day three, signal receiver cells were detached with trypsin-EDTA, and 100 µL of DMEM media containing 20,000 cells were added to tissue culture 96-well plates (Coastar). In parallel, HEK293T-expressing WT DLL4 or Delta^MAX^ were detached with trypsin-EDTA, counted manually, and dilutions prepared such that 100 µL of DMEM containing 20,000 signal sender cells were added over signal receiver cells (cell ratio 1:1). Doxycycline (Sigma) at 2 µg/mL was used to induce Notch-Gal4 expression. On day four, luciferase firefly and *Renilla* signals were determined with Dual-Glo luciferase assay system (Promega) using a GloMax luminometer (Promega). A luminescence ratio firefly:*Renilla* was determined for every well and normalized to the signal of a well containing U2OS reporter cells and HEK293T non-expressing DLL4 ligands. Notch activation is represented as the mean fold-change over the control ± S.D. of three independent biological replicates.

### Notch signaling assay using multiple ligand-presentation formats

Notch activation was assayed in different formats (Extended Data Fig. 8a) including (1) non-specific orientation (100 nM of Delta^MAX^ on well, Fig. 3a), (2) ligand-oriented coupling (10 nM of Delta^MAX^ on well, Fig. 3b), (3) magnetic beads coated with biotinylated Delta^MAX^ (ratio 1:20 reporter cells:beads, optimization of ratio is detailed in Extended Data Fig. 8d), (4) HEK293T-expressing Delta^MAX^ (ratio 1:1, signal receivers:senders, optimization of ratio can be seen in Extended Data Fig. 8c), and (5) yeast-expressing Delta^MAX^ (ratio 1:10, reporter:yeast, optimization of ratio is detailed in Extended Data Fig. 8b). On day one, U2OS hNotch1-Gal4 cells were detached with trypsin-EDTA, counted in the microscope, and 5x10E5 cells/well transferred to 6-well tissue culture plates (Falcon). On day two, U2OS cells were transfected as described above. On day three, the different activation assays were set up at the same time. For ligand oriented-coupling, streptavidin plates were coated with 10 nM of Delta^MAX^ as described above. For non-specific adsorption, non-biotinylated Delta^MAX^ at 100 nM was coated as described previously. For activation using streptavidin beads coated with biotinylated Delta^MAX^, 200 mg of streptavidin beads (streptavidin MagneSphere paramagnetic particles, Promega) were washed 3 times with 200 µL of DPBS + 0.1% BSA using a magnet (Miltenyi), and combined with 20 µg of biotinylated Delta^MAX^ for 1.5 hours at 4°C. Next, unbound Delta^MAX^ was washed out three times with 200 µL of DPBS + 0.1% BSA, and 100 µL of complexed beads (pre-diluted 1/25 in DPBS) were added to wells. For co-culture activation with HEK293T-expressing Delta^MAX^, cells were detached with trypsin-EDTA, quenched with DMEM media, washed with DPBS, and 100µL containing 20,000 cells in DMEM were added to wells. For activation with yeast cells expressing Delta^MAX^ on surface, pYAL vector encoding for Delta^MAX^ (N-EGF5) was electroporated previously into yeast cells, and DLL4 expressed in SGCAA as previously described. Delta^MAX^ expression was induced 48 hours before Notch activation assay set up. One hundred microliters of yeast cells (2x10E6 cells/mL) in serum-free DMEM were added to the assay plate. Once all the wells contained the desired activation assays, U2OS reporter cells were detached with trypsin-EDTA, and 20,000 cells in 100 µL of DMEM were added per well. All the conditions were supplemented with 2 µg/mL of doxycycline to induce Notch1-Gal4 expression. The assay plate was incubated for 24 hours at 37°C and 5% CO_2_. On day four, media was aspirated off and Luciferase Dual-Glo kit (Promega) used to detect firefly and *Renilla* signals in a luminometer (GloMax, Promega). Negative controls were BSA 0.5% for non-specific absorption, streptavidin and BSA 2% for ligand oriented-coupling, uncoated streptavidin beads for activation with magnetic beads, HEK293T non-expressing DLL4 for co-culture activation, and yeast expressing PDL-1 ECD for co-culture with yeast. The expression of Delta^MAX^ in HEK293T and yeast cells was confirmed by flow cytometry (BD Accuri C6 plus) following staining for 1 hour at 4°C with anti-hDLL4 PE antibody using DPBS + 0.5% BSA, and selection buffer, respectively (Extended Data Fig 8e). Notch activation is represented as the mean fold-change over the corresponding control ± S.D. of three independent biological replicates.

### Notch inhibition assays with soluble antagonists and WT DLL4 coated plates

For CHO-K1 reporter cells, on day one, biotinylated WT DLL4 was pre-coated in 96-well plates at 50 nM following the same protocol described above. Next, CHO cells were detached with trypsin-EDTA, washed, and diluted at 200,000 cells/mL in α-MEM media. Ten-fold serial dilutions of non-biotinylated DLL4 ligands (N-EGF5) were prepared in DPBS starting at 600 nM. Next, soluble inhibitors were mixed in ratio 1:1 with CHO-K1 200,000 cells/mL stocks for 15 min at room temperature to pre-block Notch receptors. Two hundred microliters of inhibition reactions containing 20,000 reporter cells were added to WT DLL4-coated wells, and plates incubated for 24 hours at 37°C and 5% CO_2_. When DAPT (gamma-secretase inhibitor, Sigma) and BB-94 (ADAM metalloprotease inhibitor, Sigma) were used, stock solutions were prepared at 100 µM in 8% DMSO. Appropriate DMSO controls were included to test Notch activation and cell viability. Final DMSO concentration in the experiment was kept below 1%. For comparisons of Delta^MAX^ with DAPT and BB-94, soluble inhibitors were serially diluted in DPBS using 9-folds, incubated with CHO reporter cells, and added to wells as described above. On day two, assay plates were washed with 200 µL of DPBS, CHO cells detached with trypsin-EDTA, quenched with 200 µL of α-MEM media, and Notch activation measured by flow cytometry (BD Accuri C6 Plus). The data represent the mean fluorescent intensity as percentage of Notch activation, relative to Notch activation in the absence of inhibitors ± S.D. of three independent biological replicates.

For U2OS cells, on day one, 5x10E5 cells were plated in 6-well tissue culture plates. On day two, U2OS cells were transfected as described above. On day three, 96-well plates (Coastar) were coated with streptavidin and biotinylated WT DLL4 at 50 nM as described above. Next, U2OS reporter cells were detached with trypsin-EDTA and diluted at 200,000 cells/mL in DMEM media. Ten-fold serial dilutions of soluble DLL4 variants starting at 6,000 nM were prepared in DPBS and used to pre-block reports cells following the same protocol described above. Two-hundred microliters of inhibition reactions were added to 96-well plates pre-coated with biotinylated WT DLL4 ligand at 50 nM, and assay plates incubated for 24 hours at 37°C and 5% CO_2_. On day fourth, luciferase signals were measured with Dual-Glo luciferase assay system (Promega) using a GloMax luminometer (Promega). Doxycycline at 2 µg/mL was maintained to ensure binding of soluble inhibitors to Notch-pan receptor. A ratio between firefly and *renilla* luciferase was calculated, and Notch activation normalized to wells where inhibitors were non-added. The data represent fold-change of Notch activation as percentage, relative to Notch activation in the absence of inhibitors ± S.D. of three independent biological replicates.

### Notch inhibition assay in co-culture format

For inhibition of Notch ligand-expressing cell lines, on day one, U2OS Notch3 reporter cells (signal receivers) were detached with trypsin-EDTA, washed with DPBS, and added to 6-well plates. On day two, U2OS cells were transfected as described above. On day three, signal receiver cells were detached with trypsin-EDTA, and dilutions of 400,000 cells/mL were prepared in DMEM. On the one hand, soluble DLL4 variants were prepared at 6 µM in DPBS, combined in 1:1 dilution with signal receiver cells, and incubated for 15 min at RT. On the other hand, HEK293T-expressing WT DLL4, DLL1, JAG1, JAG2 were detached with trypsin-EDTA, washed with DPBS and 200,000 cells/mL stocks prepared in DMEM. One hundred microliters of both signal senders and pre-blocked signal receivers were combined in cell ratio 1:1 and incubated for 24 hours at 37°C, and 5% CO_2_. On day four, luciferase firefly and *Renilla* signals were determined with Dual-Glo luciferase assay system (Promega) using a GloMax luminometer (Promega). Doxycycline at 2 µg/mL was kept for all the conditions. A luminescence ratio firefly:*Renilla* was determined for every well. Notch activation of each Notch3:ligand pair was normalized to the signal of Notch3 reporters in the presence of HEK293T lacking ligands. Each Notch signaling pair was set at 100% activation and soluble DLL4 treatments were relative to its corresponding pair. The data represent fold-change of Notch activation as percentage, relative to Notch activation in the absence of inhibitors ± S.D. of three independent biological replicates.

For inhibition of CHO-K1 cells with DAPT and Delta^MAX^, the experiment was set-up following the protocol described above with the exception that CHO-K1 reporters were incubated with HEK293T-expressing WT DLL4 for Notch activation. Three-fold serial dilutions of soluble Delta^MAX^ and DAPT were prepared in DPBS starting at 3 µM. For DAPT, a stock solution of 100 µM in 8% DMSO was prepared before serial dilutions in DPBS. Appropriate DMSO controls were also assayed for cytotoxicity. Final DMSO concentration in the wells was maintained below 1%. Thirty thousand signal senders and pre-blocked signal receivers (100 µL each) were incubated for 24 hours at 37°C and 5% CO_2_ before analysis by flow cytometry. Maximum Notch activation (100%) was set in the absence of soluble inhibitors and normalized to wells containing CHO-K1 cells and HEK293T lacking expression of WT DLL4. Mean fluorescence intensity is represented as percentage relative to Notch activation in the absence of inhibitors ± S.D. of three independent biological replicates.

### Co-culture assays for T-cell activation and proliferation

Functional assays for human CD8^+^ T cells were performed in co-culture with K562 cells expressing human CD32 (K32 cells) generated as described before^5^. Briefly, cells were γ-irradiated (100Gy) and washed twice with PBS before being loaded with OKT3 (0.0125-0.5μg/mL, BioXcell) antibody at RT for 10 min. Human CD8^+^ T cells were negatively isolated from PBMC and labeled with 5μM of Cell Trace Violet (Invitrogen) according to manufacturer’s specifications. Next, CD8^+^ T cells were co-cultured with K32 cells at ratio 1:10 (T cell: K32). The frequency of proliferating T cells was determined after 96 hours by flow cytometry using a CytoFLEX II (Beckman Coulter). Zombie Green (Biolegend) was used for life/dead cell staining to discard dead cells from the analysis. A set of samples were additionally incubated with Golgi stop (BD) at 0.5 μL/well for 5 hours and washed with PBS for detection of intracellular IFNγ using anti-hIFNγ APC antibody (Biolegend). The frequency of IFNγ positive cells was analyzed by flow cytometry (CytoFLEX II, Beckman Coulter). Dead cells were excluded from the analysis by staining with Zombie green (Biolegend). The data represent the mean percentage of proliferating T cells or IFNγ^+^ T cells ± S.E.M of four independent biological replicates.

### Quantitative Real-time PCR

Human CD8^+^ T cells were co-cultured with K32-DLL4 cells for 96 hours and positively enriched by MACS column (Invitrogen Kit). Total RNA was isolated from CD8^+^ T cells using TRIzol (Life Technologies). Reverse transcription reaction was performed using Verso cDNA Synthesis Kit (Thermo Scientific). Quantitative PCR reactions were prepared by using Bio-Rad SYBR green master mix and performed on an Applied Biosystems thermocycler (7900 HT). Relative expression was calculated using the ΔΔCt method and normalized to β2M levels. The data represent the mean quantity of mRNA (in-fold) for different T cell markers ± S.E.M of four independent biological replicates. Pair primers used were:

Human β2-Microglobulin

Forward: 5’-ATGAGTATGCCTGCCGTGTGA

Reverse: 5’-GGCATCTTCAAACCTCCATG

Human IFNγ

Forward: 5’-GACCAGAGCATCCAAAAGAG

Reverse: 5’-GGACATTCAAGTCAGTTACCGAATA

Human Granzyme B

Forward: 5’-CCCTGGGAAAACACTCACACA

Reverse: 5’-GCACAACTCAATGGTACTGTCG)

Human Hes-4

Forward: 5’-ACGGTCATCTCCAGGATGT

Reverse: 5’-CGAGCGCGTATTAACGAGAG)

### Immunoblot Analysis

For detection of Notch activation in CD8^+^ T cells, following co-culture with K32 cell lines, CD8^+^ T cells were positively enriched by Magnisort Human CD8^+^T cell positive selection Kit (Invitrogen). Nuclear fraction was isolated using NE-PER Nuclear and Cytoplasmic extraction kit (ThermoFisher). Equal protein amounts of nuclear cell lysates were run on 4-20% gradient Tris-glycine gels (Novex-Invitrogen), transferred to PVDF membranes using an iBlot Gel Transfer Device (ThermoFisher), and immunoblotted with antibodies anti-cleaved Notch1 (Val1744 rabbit mAb, Cell Signaling Technology, 1:1,000), anti-Notch2 (D76A6 rabbit mAb, Cell Signaling Technology, 1:1,000), and anti-β Actin (mouse mAb, Sigma, 1:20,000). Secondary antibodies anti-IgG (anti-rabbit or anti-mouse, GE Healthcare, 1:5,000) conjugated to horseradish peroxidase (HRP) were used for detection of proteins using ECL Western Blot Substrate Reagent (Thermo Fisher Scientific). Images were acquired using a Chemidoc Imaging System and analyzed with Image-Lab software (Bio-Rad).

For Notch activation in U2OS and MCF-7 cells, WT DLL4 and Delta^MAX^ biotinylated ligands were immobilized at 50 nM in 6-well streptavidin-coated plates following a scaled up protocol of the one described before. After 24 hours of stimulation, cells were lysed in Laemmli serum buffer (40% glycerol, 4% SDS, 250 mM Tris-HCl, 0.02% bromophenol blue, and 20% β-mercaptoethanol). Cell lysates were heated at 100°C for 5 min, centrifuged at 21,000*xg* for 10 min at 4°C, and supernatants stored at -20°C. Proteins were separated by SDS-PAGE (12% Mini-PROTEAN TGX Precast Protein Gels, Bio-Rad) and transferred to PVDF membranes using an iBlot2 Gel Transfer Device (Thermo Fisher Scientific). The membranes were blocked in 3% BSA + 0.1% TBS-Tween. Primary antibodies were anti-cleaved Notch1 (Val1744 rabbit mAb, Cell Signaling Technology, 1:1,000), and β-Actin (rabbit pAb, Cell Signaling Technology, 1:1,000). Secondary antibody anti-Rabbit IgG conjugated to HRP (Goat mAb, Vector Laboratories, 1:8,000) was used for detection of proteins using SuperSignal West Pico PLUS Chemiluminescent Substrate (Thermo Fisher Scientific). Images were acquired using an Amersham Imager 600 (GE Healthcare), and contrast adjusted with a curves layer.

## EXTENDED DATA FIGURES

**Extended Data Fig 1.**
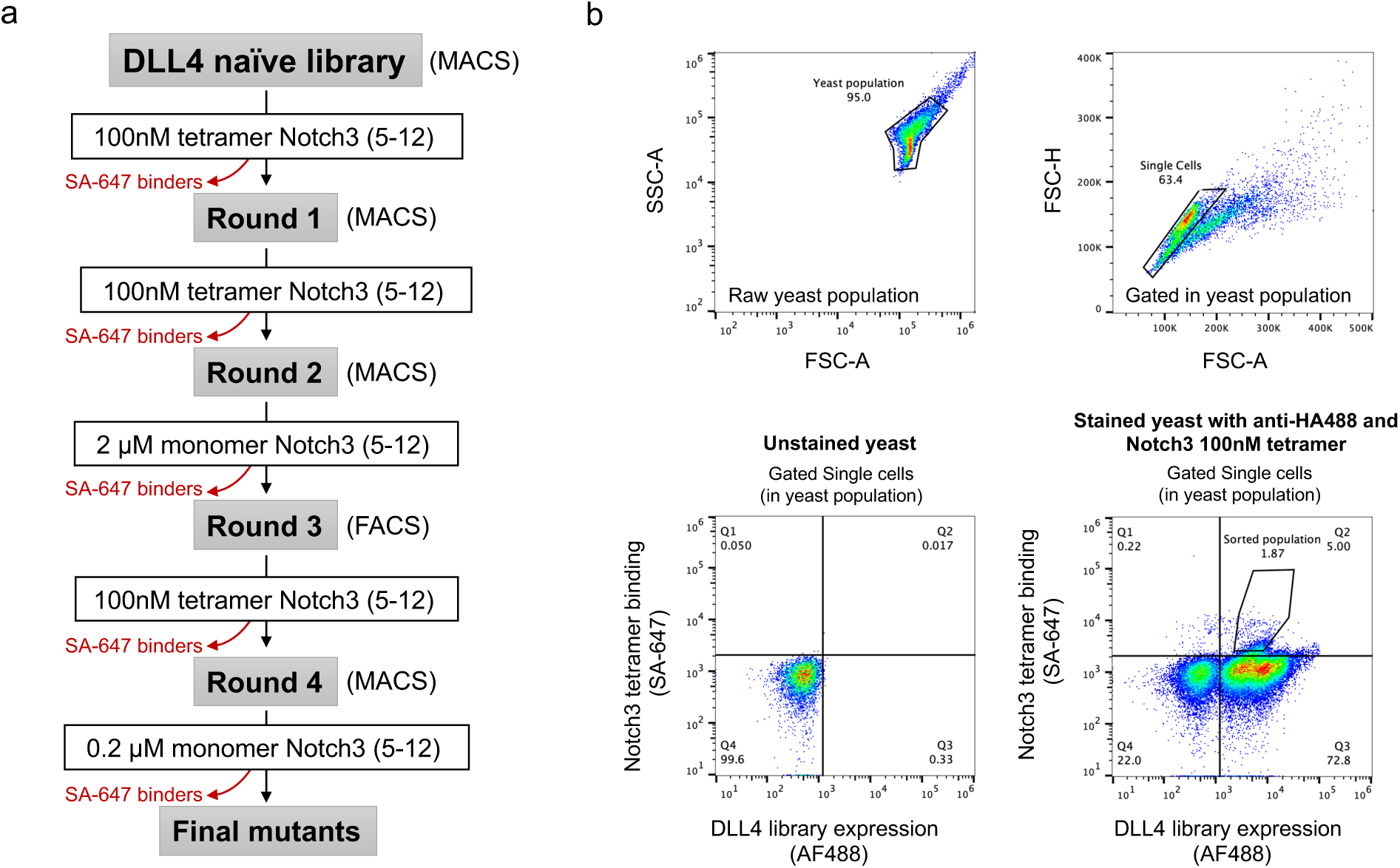
Selection strategy for isolation of high-affinity DLL4 populations. (**A**) Flow chart depicting the selection strategy used to isolate high-affinity DLL4 variants. Red arrows and text indicate negative selections. (**B**) Gating strategy for sorting high-affinity binders to Notch3.

**Extended Data Fig 2.**
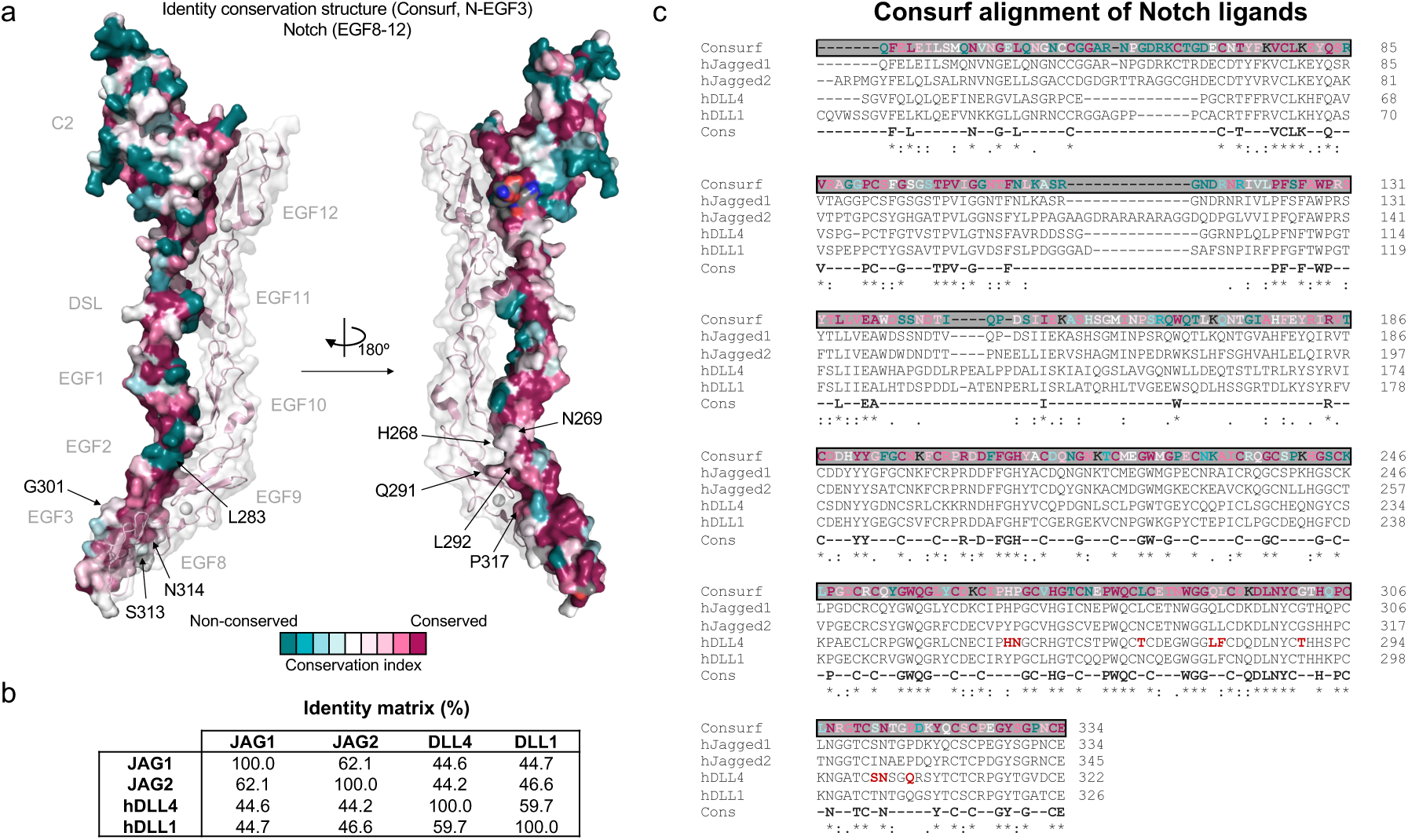
Conservation analysis of Site 3 library positions in activating Notch ligands. (**A**) A structural model depicting amino acid conservation in DLL1, DLL4, Jag1, and Jag2 residues was generated in Consurf using rat JAG1 as a template (PDB ID: 5UK5). (**B**) Identity matrix indicating the sequence identity between human Notch ligands. (**C**) Sequence alignment depicting conservation at each residue position. Residues mutated in the DLL4 mutant library are highlighted in red.

**Extended Data Fig 3.**
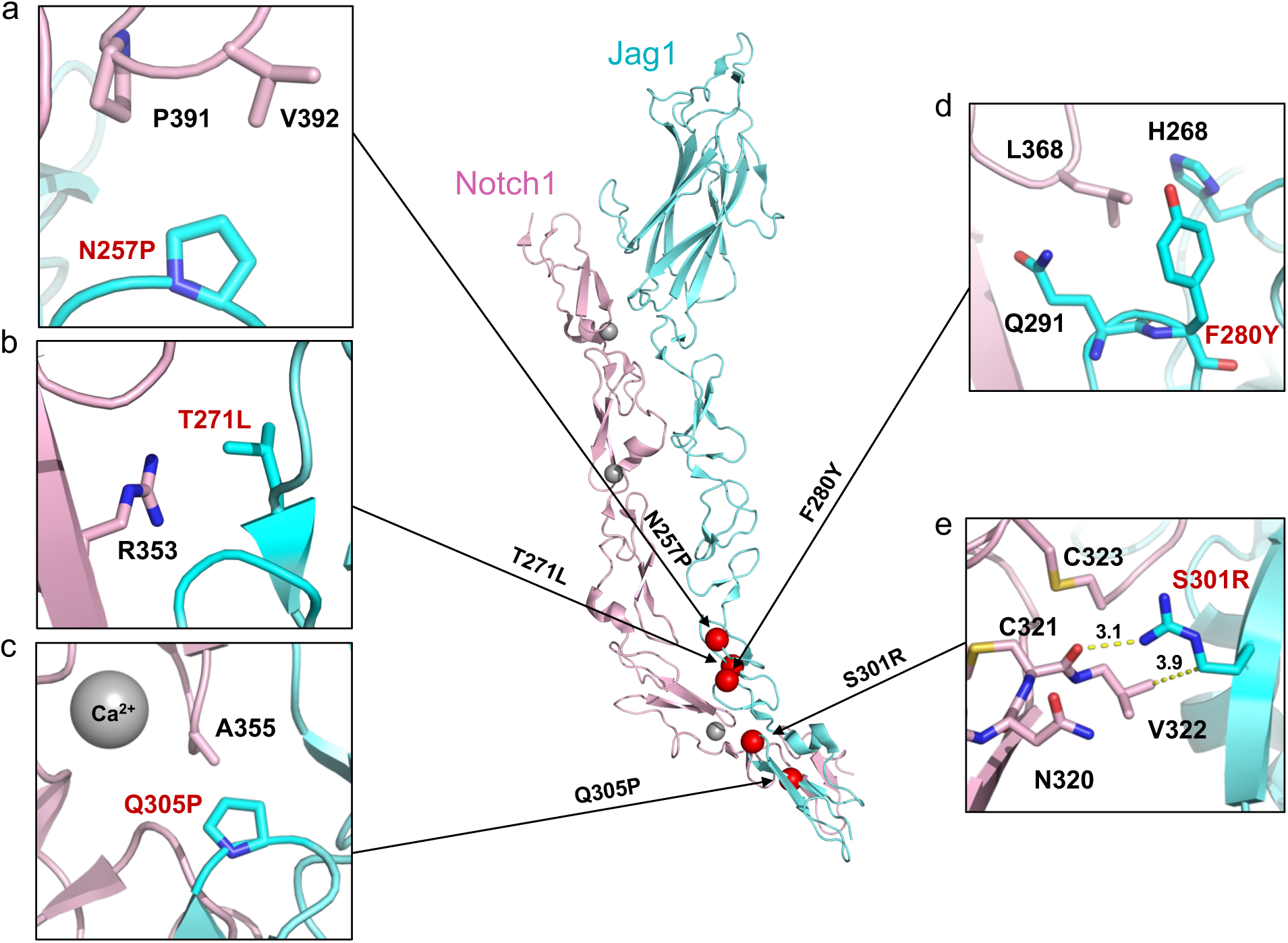
Mutational analysis of evolved DLL4 Site 3 over Jagged1:Notch1 structure. Cartoon representation showing the structural context of DLL4.v2 mutants in a model of the rat Jag1-Notch1 complex (PDB ID:5UK5). (**A**), (**B**), (**C**), (**D**), and (**E**) are zoom panels depicting the residues that surround the mutated position. Numbers and dashes in **(E)** are inter-atomic distances atoms measured in angstroms.

**Extended Data Fig 4.**
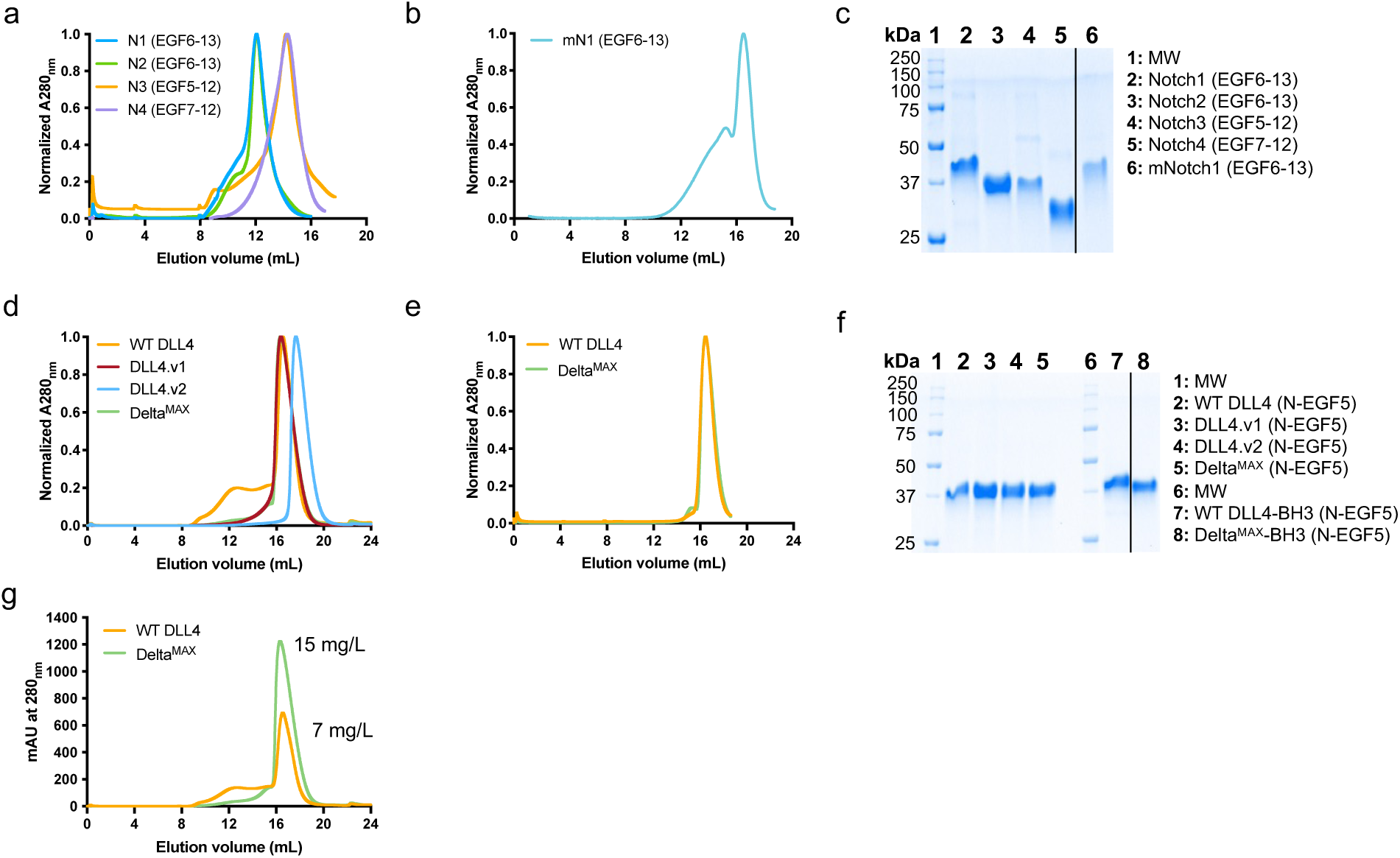
Purification of Notch and DLL4 fragments for SPR studies and signaling assays. **(A-B)** SEC chromatograms from the purification of the ligand-binding regions of human Notch1-4 **(A)** and murine Notch1 **(B)**. All proteins were purified by Ni-NTA followed by size exclusion chromatography using a Superdex S75 column. **(C)** SDS-PAGE gels showing the purity and molecular weight of each Notch construct. (**D**) SEC chromatograms from the purification of DLL4, DLL4.v1, DLL4.v2, and Delta^MAX^ proteins. All proteins were purified by Ni-NTA followed by size exclusion chromatography using a Superdex S200 column. **(E)** SEC chromatograms from the purification of C-termini biotinylated WT DLL4 and Delta^MAX^ proteins. **(F)** SDS-PAGE gels showing the purity and molecular weight of each DLL4 construct. **(G)** SEC chromatograms from 1L protein preps were overlaid to highlight the increased yield of recombinant Delta^MAX^ compared to WT DLL4.

**Extended Data Fig 5.**
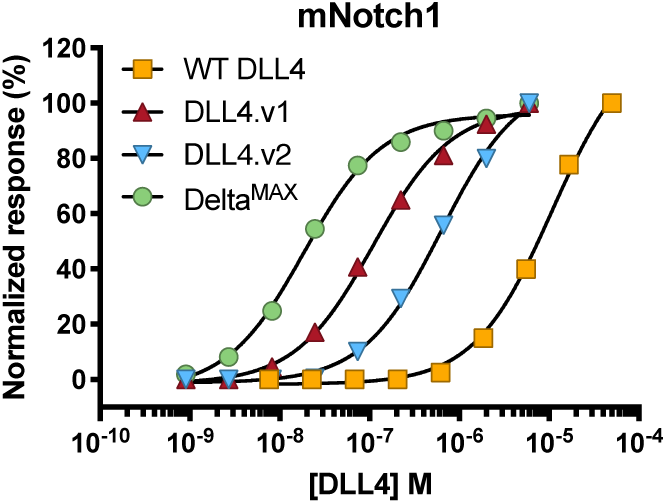
SPR binding assays for mNotch1. SPR binding isotherms were fitted to a 1:1 binding model to determine the binding affinity between mNotch1(6-13) and each DLL4 variant.

**Extended Data Fig 6.**
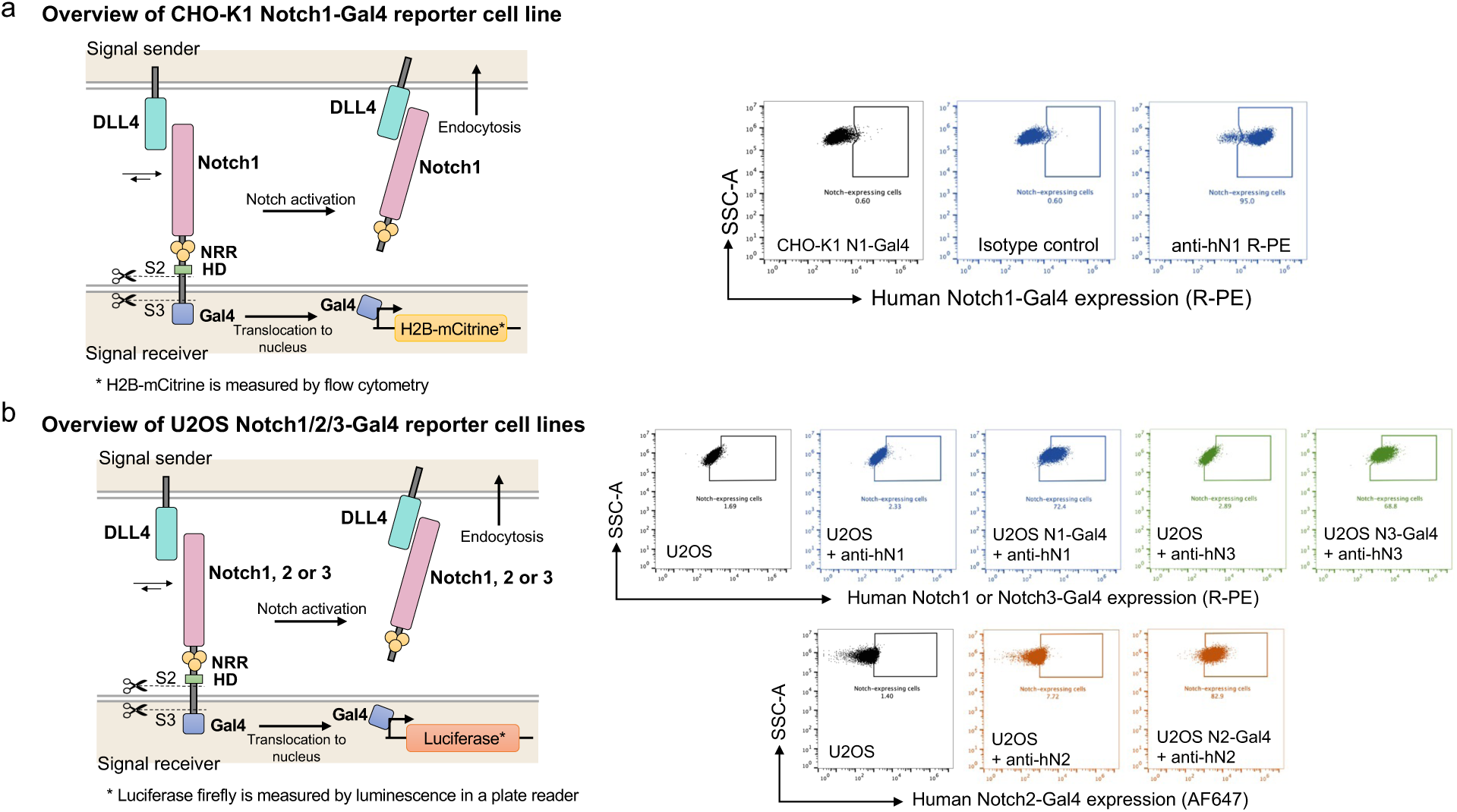
Notch reporter cell lines used in this study. Cartoon schematics describing the fluorescent **(A)** and luminescent **(B)** Notch-Gal4 reporter systems used for signaling assays. Flow cytometry dot plots depict the staining of each cell line with Notch-specific antibodies to detect surface expression.

**Extended Data Fig 7.**
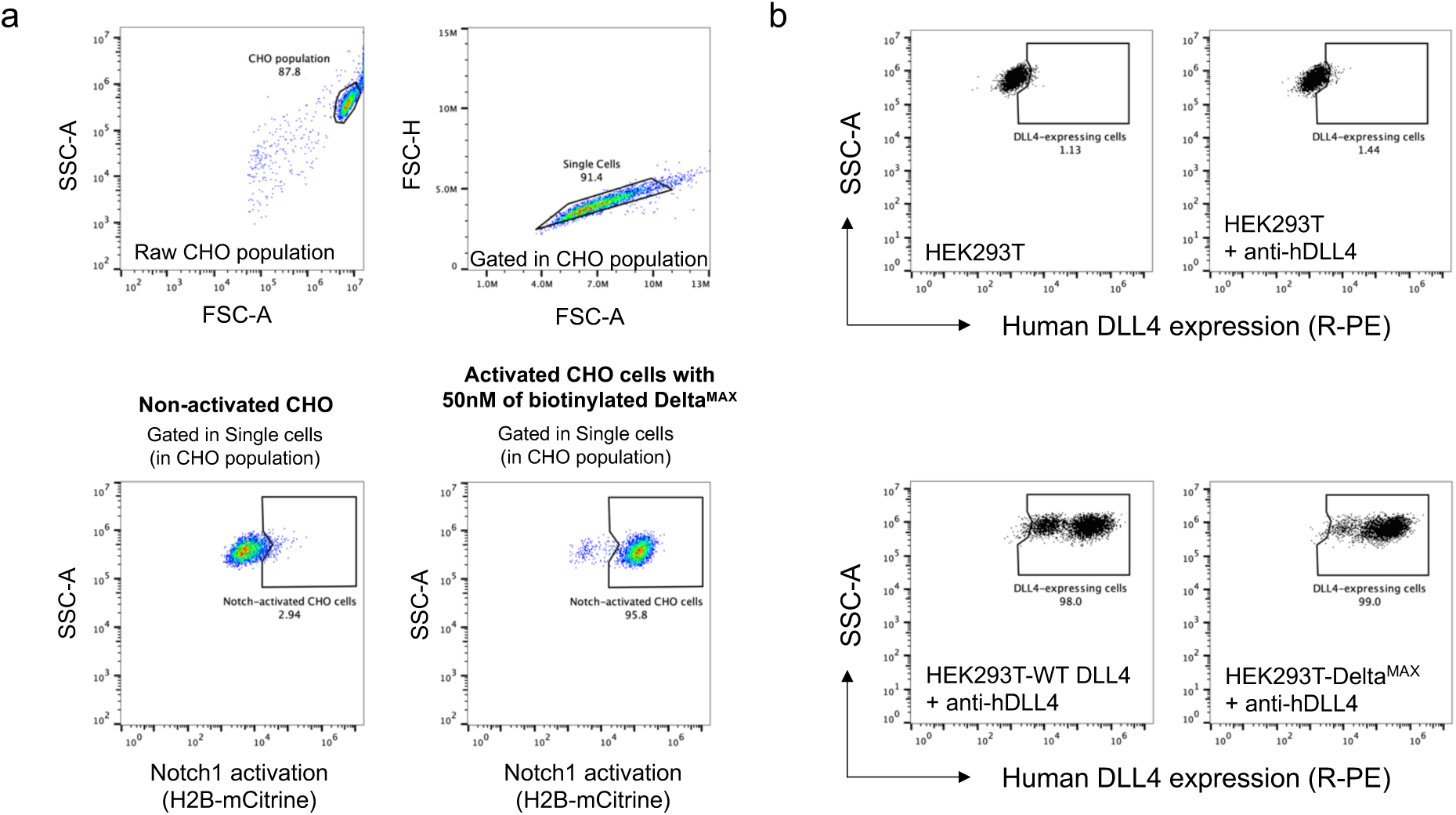
Gating strategy for Notch activation assays using CHO-K1 and detection of DLL4 expression in HEK293T stable cell lines. (**A**) DLL4 variants were non-specifically adsorbed to 96-well tissue culture plates or immobilized to streptavidin plates. Next, CHO-K1 Notch1-Gal4 cells were added to plates and Notch activation was measured by flow cytometry. The gating strategy to quantify Notch1 activation based on expression of H2B-mCitrine using CHO reporter cells is detailed. HEK293T were transduced to generate stable cell lines expressing WT DLL4 or Delta^MAX^ and sorted to normalize the expression of DLL4. (**B**) Expression of WT DLL4 and Delta^MAX^ on HEK293T was analyzed with anti-hDLL4 PE antibody using flow cytometry. The sorted cell lines were used for Notch activation assays in co-culture with Notch1, Notch2, and Notch3-U2OS luciferase reporters.

**Extended Data Fig 8.**
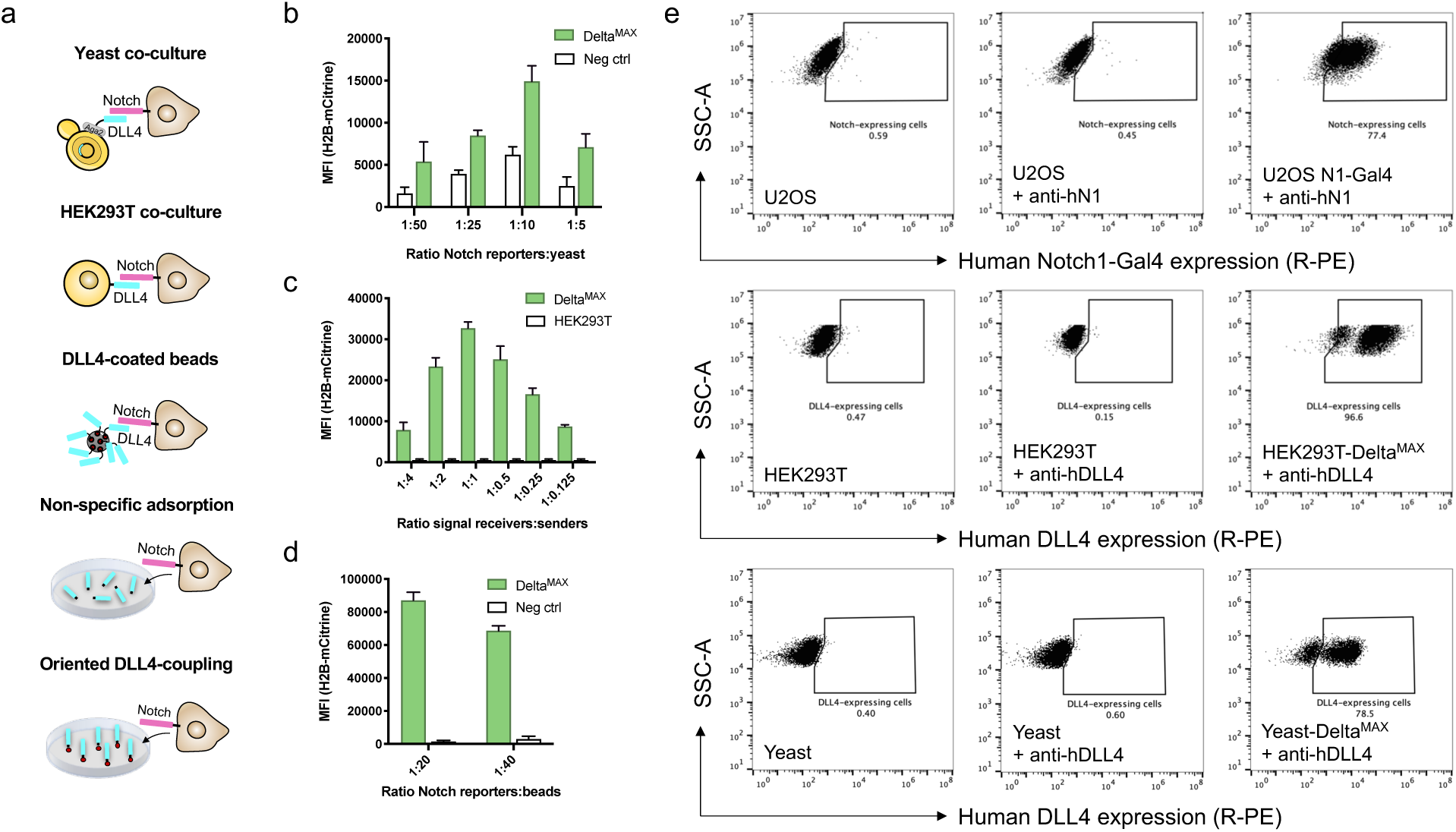
Optimization for all-in-one assay using Delta^MAX^. (**A**) Cartoon representation of different ligand-presentation formats. (**B**) Yeast expressing Delta^MAX^ (N-EGF5) were co-cultured with CHO-K1 reporter cells for 24h at different ratios and fluorescence was measured by flow cytometry. (**C**) HEK293T cells expressing Delta^MAX^ were co-cultured with Notch1 reporter cells at various ratios for 24h and Notch activation was measured by flow cytometry. (**D**) Magnetic beads were pre-coated with biotinylated Delta^MAX^ and co-cultured with CHO K1 reporter cells at ratios of 1:20 or 1:40 to stimulate Notch activation. (**E**) Flow cytometry dot plots depict the expression level of Notch1 and Delta^MAX^.

**Extended Data Fig 9.**
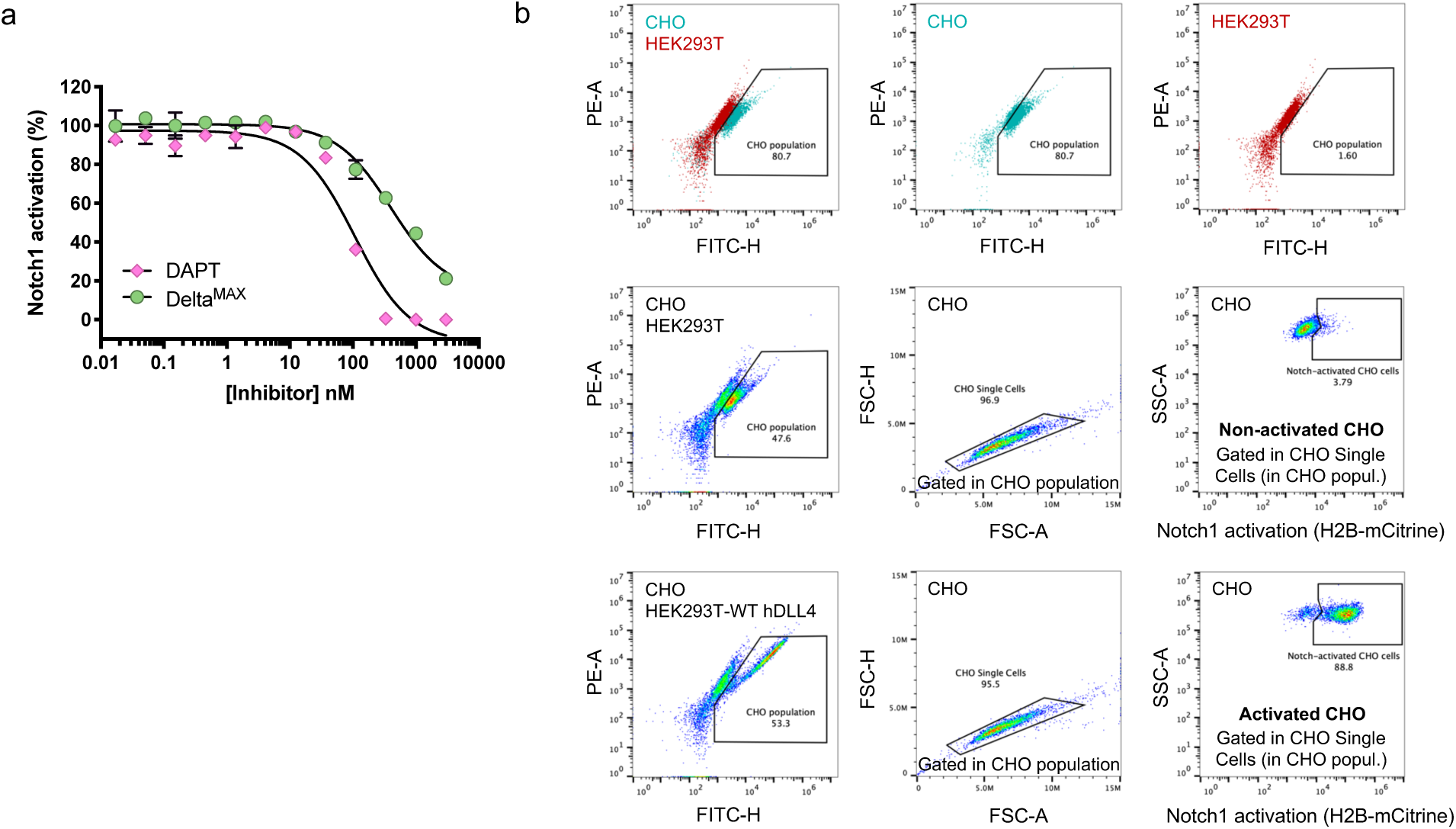
Co-culture inhibition using CHO-K1 cells, DAPT, or Delta^MAX^ as soluble antagonists. (**A**) Dose-titration assay comparing the inhibition potency of DAPT and Delta^MAX^. Fluorescent Notch1 reporter cells were cultured in a 1:1 ratio with HEK293T cells stably expressing WT DLL4. (**B**) Summary of the gating strategy used for flow cytometry to differentiate between HEK293T and CHO cell signals. The basal expression of H2B-mCitrine in CHO cells was used as a criterion to distinguish between cell types. Notably, the population identified in the middle and bottom panels (Notch1 reporter CHO cells) corresponds to approximately 50% of the total cells, which is consistent with a 1:1 ratio of CHO cells to 293T cells.

**Extended Data Fig 10.**
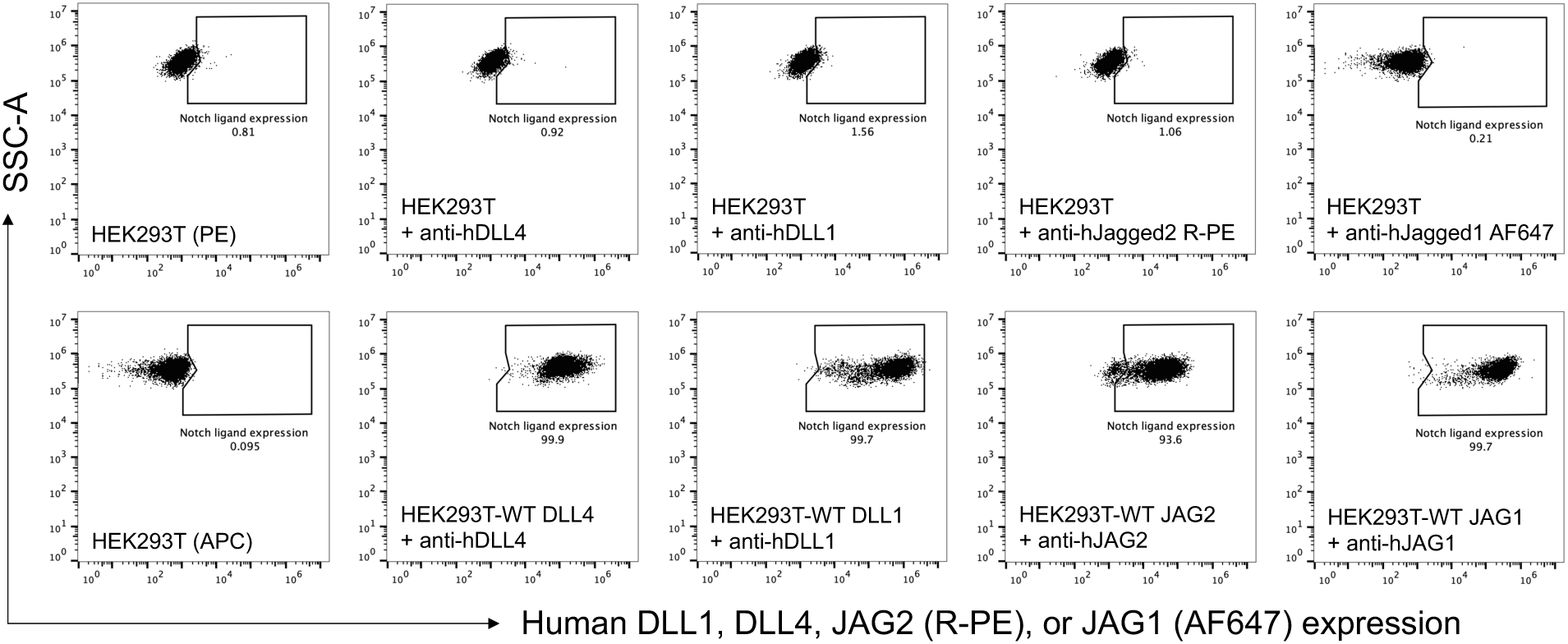
Detection of Notch ligands expressed on HEK293T stable cell lines by flow cytometry. HEK293T stable cells lines expressing WT DLL4, DLL1, JAG1, or JAG2 were stained with specific antibodies targeting the ECDs of each Notch ligand and measured by flow cytometry. The expression of Notch ligands, as well as sequencing results, validated these stable cell lines.

